# Putative APOBEC3 deaminase editing in MPXV as evidence for sustained human transmission since at least 2016

**DOI:** 10.1101/2023.01.23.525187

**Authors:** Áine O’Toole, Richard A. Neher, Nnaemeka Ndodo, Vitor Borges, Ben Gannon, João Paulo Gomes, Natalie Groves, David J King, Daniel Maloney, Philippe Lemey, Kuiama Lewandowski, Nicholas Loman, Richard Myers, Marc A. Suchard, Michael Worobey, Meera Chand, Chikwe Ihekweazu, David Ulaeto, Ifedayo Adetifa, Andrew Rambaut

## Abstract

Mpox is often described as being endemic in West and Central Africa as a zoonotic disease that transmits through contact with the reservoir rodent host, likely a species of African squirrel. In May 2022, human cases of Mpox were detected spreading internationally beyond countries with known endemic reservoirs. At time of writing, 84,700 confirmed cases have been reported in 110 countries. When the first cases from 2022 were sequenced, it was seen that they shared 42 single nucleotide differences from the closest mpox virus (MPXV) genome sampled in 2018. This number of changes within 3-4 years is unexpectedly large and points to a much greater evolutionary rate than expected for a poxvirus. Strikingly, most nucleotide changes are of a specific type – a dinucleotide change from TC->TT or its reverse complement GA->AA. This mutation type is characteristic of the action of APOBEC3 deaminases; host-enzymes with reported antiviral function. Analysis of MPXV genomes sampled from 2017 to 2022 showed further evidence of TC->TT mutation pattern enrichment, with 93% of transmitted single nucleotide mutations since 2017 consistent with APOBEC3 editing. Assuming APOBEC-editing is characteristic of MPXV infection in human hosts, we propose an APOBEC clock that – at a rate of ~6 APOBEC3 mutations per year – estimates MPXV has been circulating in humans since 2016. This evolutionary pattern of host-enzyme editing has implications for the longer-term fitness of the virus in this epidemic as such mechanisms are primarily antiviral in function, but in the context of a poxvirus also provide a source of variation that may conceivably facilitate adaptation.

## Introduction

Since 2017, the Nigeria Centre for Disease Control has been reporting cases of MPXV (mpox virus) infection in humans (*1*). MPXV is often described as being endemic in West and Central Africa as a zoonotic disease that transmits through contact with the reservoir rodent host. The virus is divided into two clades with isolates from Central Africa falling into Clade I and isolates from West Africa predominantly falling into Clade II. Whilst MPXV has a broad host range and can readily infect multiple rodent and primate species in the laboratory and in nature (*2–8*), African rodents remain the most likely reservoir hosts, specifically rope and sun squirrels (genera *Funisciurus* and *Heliosciurus*), African dormice (genus *Graphiurus*) and pouched rats (genus *Cricetomys*), native to West and Central Africa. MPXV-specific antibodies have been detected in squirrel species in a number of surveys that sampled wildlife from areas of reported human MPXV cases (*3–5*) and in 1985 an infected rope squirrel was associated with a zoonotic outbreak in the Democratic Republic of Congo (*9*). Pouched rats and dormice housed in proximity to North American prairie dogs were the likely cause of a widespread zoonotic MPXV outbreak in the USA in 2003 (*10*). Ecological niche modelling predicted a dramatic range expansion of the rope squirrel with climate change by 2050, however this is not common to all putative host species as the range of the pouched rat (*Cricetomys gambianus*) is predicted to contract in response to climate change (*11*). Furthermore, continued forest clearing in favour of agricultural fields and forests may actually increase habitat suitable for reservoir hosts, such as the rope squirrel, and precipitate more frequent human-reservoir interactions (*3, 11*).

Since the first human cases were observed in the 1970s MPXV infections have been predominantly associated with infants and children, with a marked increase in case counts since the 1980s, likely due in part to the end of smallpox vaccination after eradication (*12–14*). The smallpox vaccine offers protection against MPXV, however with the eradication of variola virus in 1977, protection has been waning in the population. Prior to 2017, the majority of cases have been reported in DRC, as zoonoses initiating in school-age boys with Clade I viruses (*12*). The outbreak observed in Nigeria from 2017, from which the global outbreak has grown, has involved Clade II viruses (*8, 15*). The demographic mainly affected in Nigeria has been men aged 25 to 40, with only ~25% of cases involving women, and very few cases in children (*8*).

Between 2018 and March 2022 there have been a limited number of cases in Israel (MN648051), Singapore (MT250197), USA (ON676708), and the United Kingdom (UK; MT903344, MT903343, OL504741) associated with travel from Nigeria, with one case of onward transmission in the UK (MT903345). However, in May 2022, cases of MPXV infection were detected spreading widely across Europe and subsequently across the globe. At time of writing, 84,700 confirmed cases have been reported in 110 countries (*16*), predominantly in men who have sex with men (MSM) sexual contact networks. The rapid growth of cases in these networks has prompted concern that the 2022 epidemic represents a transition to a more transmissible form of the virus.

In the wake of the SARS-CoV-2 pandemic, the expanded capacity to perform viral genome sequencing enabled rapid generation of MPXV genome data, and the legacy of open-data sharing practices led to their quick dissemination to public repositories. The first MPXV genome sequences from 2022 cases showed these viruses had descended from the clade sampled in 2017-2019 from cases diagnosed in Israel, Nigeria, Singapore, and the UK (*17*). These early 2022 genomes are indicated as a triangle within Clade IIb in Figure 1a and represent lineage B.1 as per the nomenclature proposed by Happi et al. (*15*). Isidro et al., (*17*) noticed that sequences within lineage B.1 shared 42 single nucleotide differences from the closest earlier MPXV genomes from 2018 (specifically UK_P2 and UK_P3; accession numbers: MT903344, MT903345). Later, an MPXV sample from 2021 was retrospectively sequenced (Accession number: ON676708) (*18*) and split this long branch to lineage B.1 (Figure 4).

**Figure 1.**
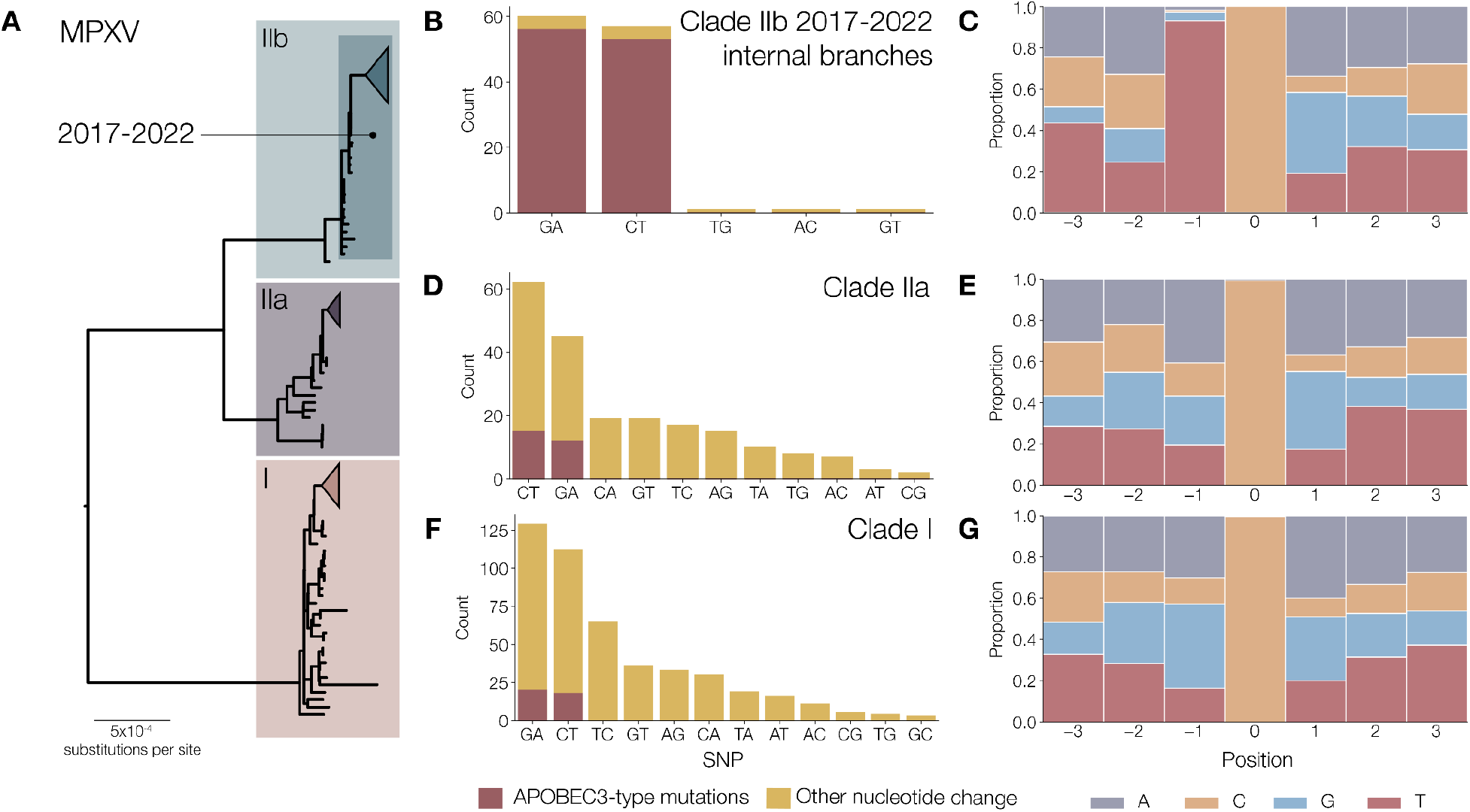
A) MPXV phylogenetic tree highlighting Clade I (predominantly sequences from the DRC), Clade IIa (predominantly West African sequences) and Clade IIb. Within Clade IIb is a sub-clade of genomes sampled from 2017-2022 that show distinct mutational patterns to the other two clades. B) Of 120 reconstructed single nucleotide mutations that occurred on internal branches of the Clade IIb phylogeny (so are observed transmitted mutations), 109 are consistent with APOBEC3 editing (90.8% of mutations). C) Observed heptamers of C->T or G->A mutated sites in internal branches of the IIb phylogeny. Heptamers associated with G->A mutations have been reverse-complemented to reflect deamination on the negative strand. Most C->T mutations are present in a TC dimer context, consistent with APOBEC3 editing (107 of 115 mutations, or 93%). D) Ancestral state reconstruction performed across Clade IIa does not produce the same enrichment of mutations consistent with APOBEC3 editing, with only 27 of 207 observed mutations (13%) fitting the dinucleotide pattern. E) Observed heptamers around C->T mutations (and reverse-complemented G->A mutations) in the Clade IIa phylogeny. 29 of 149 (19%) mutations have the dinucleotide context of APOBEC3. F) Only 38 of 463 Clade I mutations (8%) are consistent with APOBEC3 editing. G) Observed heptamers in C->T mutations (and reverse-complemented G->A mutations) in the Clade I phylogeny respectively. 42 of 256 (16%) normalised C->T or G->A mutations are in a TC or GA dinucleotide context, which is what we would predict under standard models of nucleotide evolution.

**Figure 2.**
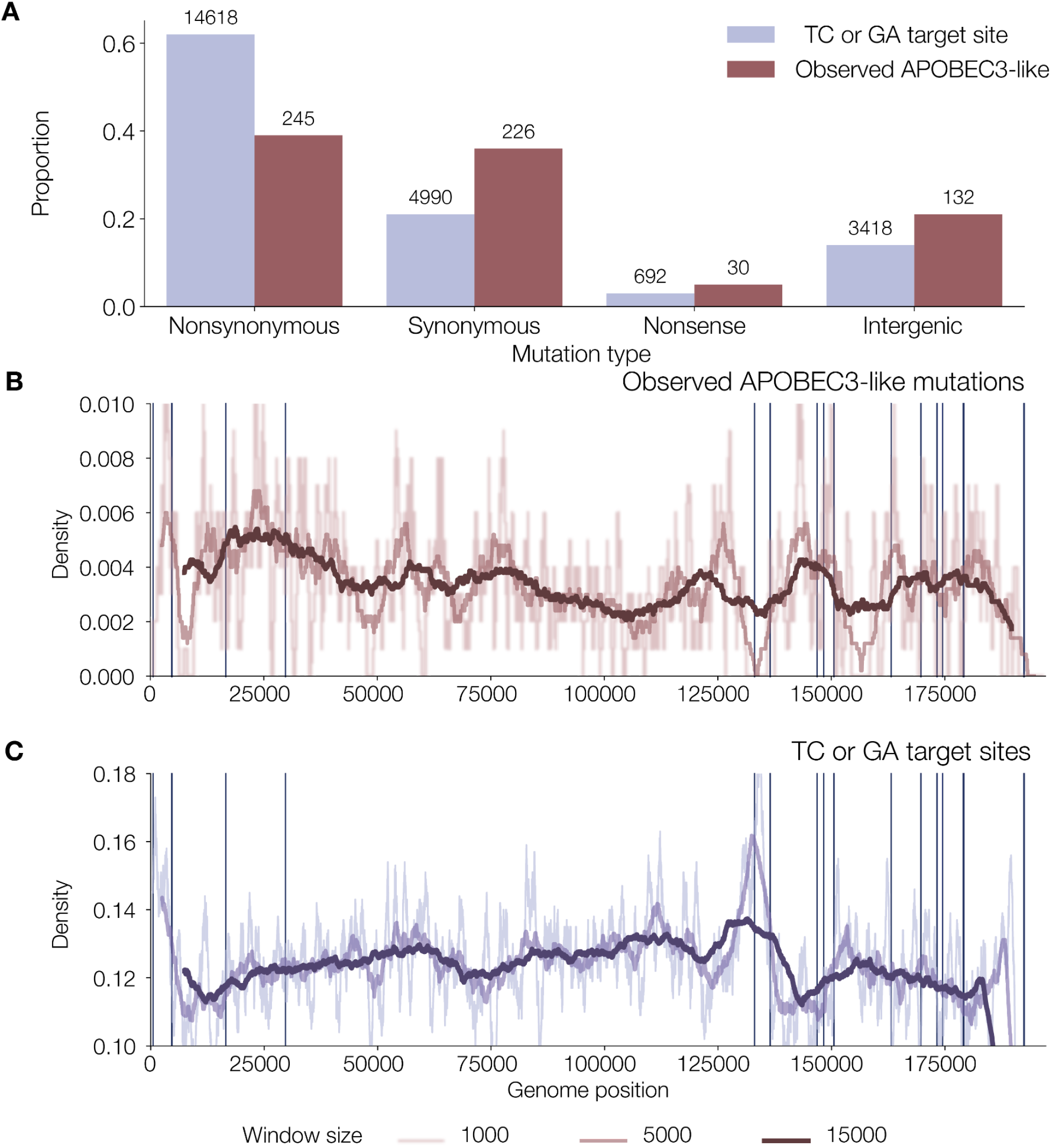
A) Consequence of hypothetical APOBEC3 mutations at target dimer site (either the C in the TC target site or G in the GA target site) in currently unmutated sites in the coding regions of the NCBI reference MPXV genome (Accession NC_063383) and those observed APOBEC3 mutations across the coding regions of the Clade IIb phylogeny (including genomes in Figure 4 and Supplementary Figure 1, except the outgroup branch leading to the 1971 genome sequence). These are categorised into non-synonymous (altered amino acid), synonymous (amino acid remaining unchanged), nonsense (editing producing a stop codon) and intergenic (not present in a coding sequence). B) The density (moving average linear convolution with window sizes of 1000, 5000 and 15000) of observed APOBEC-consistent mutations and C) APOBEC-target dimer sites (GA and TC) in a masked Clade IIb reference genome (Accession NC_063383). Repetitive/ low complexity masked regions in the alignment indicated by vertical bars.

**Figure 3.**
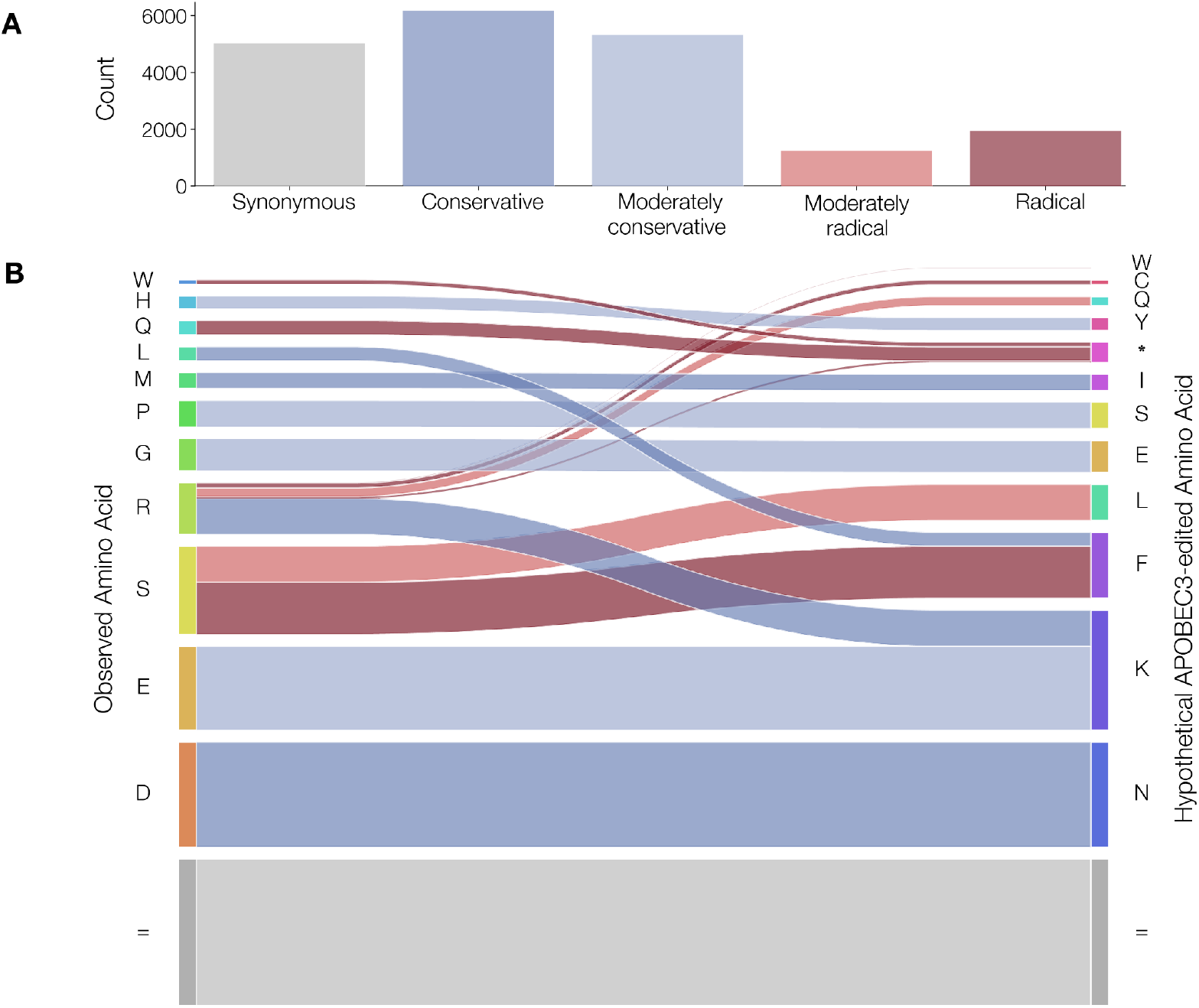
Amino acid mutations at TC and GA dimer sites in a reference MPXV genome (accession number NC_063383) that could occur through APOBEC3 editing. A) Barplot of amino acid changes categorised by Grantham Score (0-50 conservative, 51-100 moderately conservative, 101-150 moderately radical, >150 radical). B) Hypothetical amino acid changes for codons overlapping with TC and GA target sites if APOBEC3 edited those dimers to TT and AA.

**Figure 4.**
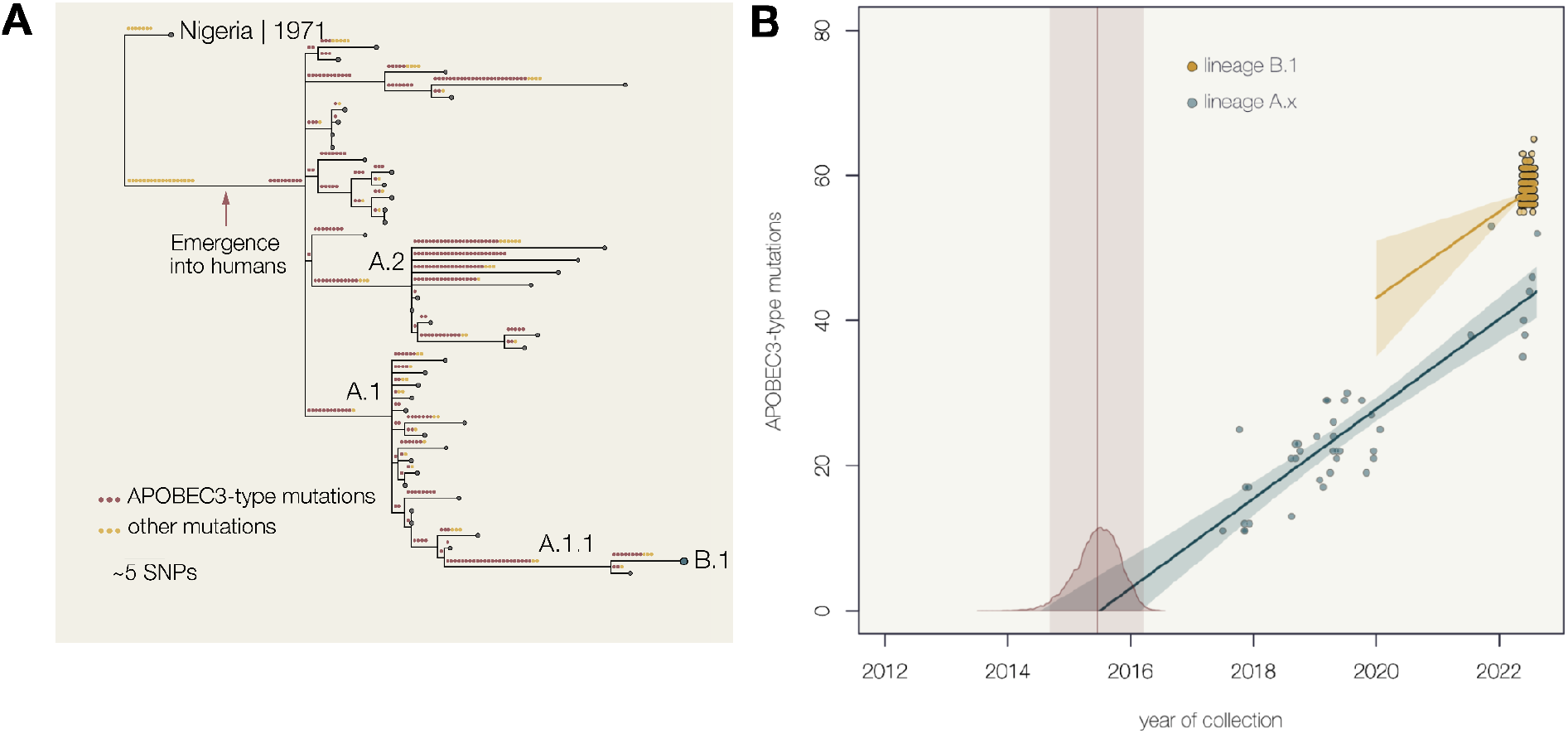
A) A phylogenetic tree of MPXV genomes sampled from human infections from 2017-2022, with an outgroup sequence from an outbreak in Nigeria in 1971 (n=45 including outgroups). Lineages indicated as per nomenclature proposed by Happi et al., (*15*). Single nucleotide mutations along each branch are indicated with circles coloured by whether a mutation is putatively APOBEC3 edited (TC->TT and GA->AA; red) or whether it is another mutation type (yellow). B) Root to tip plot of APOBEC3 mutations from MPXV genomes sampled since 2017. The reconstructed most recent common ancestor (MRCA) of the panel A phylogeny is used as the root. We performed Bayesian regression analysis on sequences from panel A, which includes one representative B.1 genome, and also separately on the B.1 lineage using the reconstructed MRCA (B.1 phylogeny shown in Supplementary Figure 1).

The long term evolutionary rate of the related variola virus (VARV; the smallpox virus) has previously been estimated to be about 9×10^-6^ (posterior mean, with 95% Bayesian credible intervals of 7.8 – 10.2×10^-6^) substitutions per site per year (*19*) translating into about 1-2 nucleotide changes per year for a nearly 200,000 nucleotide genome. During a 2017 outbreak of MPXV in chimpanzees, the evolutionary rate of MPXV was estimated to be 1.9×10^-6^ substitutions per site per year (1.2 – 2.7×10^-6^), on the same order of magnitude as the VARV estimate (~1 nucleotide change every 3 years) (*20*). This makes 42 substitutions in the space of 3-4 years an unexpectedly large number. Furthermore, across the phylogeny of 45 MPXV genome sequences collected since 2017 (Figure 4A) and within the 769 genomes available within lineage B.1 (Supplementary Figure 1), we see accumulation of an excess of mutations since 2017. If MPXV is a zoonotic virus with limited human to human transmission, this long branch could be evidence of adaptation to humans facilitating the sustained transmission that is now observed.

Strikingly, the majority of observed nucleotide changes appear to be of a particular type, a dinucleotide change from TC->TT or its reverse complement GA->AA (*21*). This specific mutation is characteristic of the action of the APOBEC3 (Apolipoprotein B mRNA editing enzyme, catalytic polypeptide 3) family of cytosine deaminases. These act on single stranded DNA (ssDNA) to deaminate cytosine to uracil causing a G->A mutation in the complementary strand when it is synthesised. Most human APOBEC3 molecules have a strong bias towards deaminating 5’TC dinucleotides, with the exception being APOBEC3G which prefers 5’CC dinucleotides (*22*). Recent work also highlighted the importance of ssDNA secondary structure flanking the target site for APOBEC3 editing (*23*).

APOBEC-type enzymes are a large and diverse family of deaminase proteins with the ability to bind RNA and ssDNA. Reported functions range from tissue-specific RNA diversification with APOBEC1, somatic hypermutation in the adaptive immune system with Activation-Induced cytidine Deaminase (AID) (*24–26*), skeletal and muscle cell differentiation with APOBEC2, and restriction of mobile elements and antiviral action in hosts with APOBEC3-type enzymes (*24, 27*). APOBEC3-driven deamination has been demonstrated with many DNA viruses and retroviruses; (*28–35*). Vaccinia virus (VACV), an Orthopoxvirus closely related to MPXV, was subjected to ABOBEC3 *in vitro* and reportedly showed no growth disadvantage (*36*). Although this study did not involve confirmatory sequencing to confirm that no APOBEC3 editing had taken place.

Whether a virus is subject to APOBEC3-induced mutation will be a function of the availability of appropriate viral target ssDNA molecules during virus replication, co-localisation of virus replication with APOBEC3, and sufficient stoichiometry of enzyme to virus genetic material for an observable amount of editing to occur but insufficient to control the virus completely. Poxviruses are large DNA viruses with complex replication cycles that transiently expose ssDNA during replication. They replicate exclusively in the cytoplasm (*37, 38*) where a number of members of the APOBEC3 enzyme family localise in addition to many other host antiviral factors. Poxviruses have evolved various antiviral factor antagonists and some have dedicated up to 50% of their genomic capacity to immune modulatory mechanisms (*39, 40*). For observable levels of APOBEC3 editing to occur, it may come down to a balance between the level of host enzyme readily available to bind target virus ssDNA and the rate of virus DNA amplification during replication.

APOBEC3 genes emerged in placental mammals from a duplication of the ancestral AID gene and have a dynamic recent evolutionary past, with gene duplication and loss across phyla (*41, 42*). APOBEC3 genes in primates have undergone recent expansion, with primate genomes now having 7 paralogs of 3 ancestral genes (*27, 43*). Rodents, the reservoir of MPXV, have only a single functional APOBEC3 protein resulting from gene loss and fusion events (*27*). This is thought to be reflected in the decreased mobile element expansion in primates relative to rodents (*44, 45*). Rodent APOBEC3 has been shown to be expressed preferentially in spleen and bone marrow with limited expression observed in other tissues (*46, 47*).

## Nomenclature

A recently published paper (*15*) proposed an updated, systematic, nomenclature for the phylogenetic structure of MPXV to replace previous geographical clade names. In this nomenclature the 2022 global epidemic lies in Clade IIb with all genomically characterised cases since 2017 designated with lineage labels similar to SARS-CoV-2. The recent human epidemic, provisionally named hMPXV-1 by Happi *et al*. (*15*), comprises a hierarchy of lineages starting with ‘A’ currently represented by genomes from 2017 cases in Nigeria. The bulk of 2022 genomes are part of a lineage denoted ‘B.1’ but other lineages (‘A.2’ and ‘A.3’) have also been reported in the USA, UK and Portugal, in most cases in individuals with international travel history. We follow this nomenclature here.

## Methods

We compiled high-quality MPXV genomes from human and non-human animal outbreaks sampled from as early as 1965 and up to the current 2022 outbreak. The dataset consisted of 94 genomes with representatives from Clades I, IIa and IIb. A second dataset comprises all Clade IIb genomes from 2017-2022 but with a single representative of lineage B.1 of hMPXV-1 and two earlier sequences from Nigeria from 1971 and 1978 (accession numbers KJ642617 and KJ642615, respectively). The genome from 1978 (accession KJ642615) was used as an outgroup, due to its greater divergence despite being sampled more recently, used to root the tree and then not analysed further. A separate analysis was run specifically on lineage B.1 where all high-quality B.1 genomes with an exact date of collection on Genbank were downloaded on the 2022-08-22. The 2021 genome from Maryland, USA (lineage A.1.1; accession number: ON676708; Ref (*18*) was used as an outgroup to root the B.1 tree (n=769). Accession numbers and acknowledgments for genome sequences used in this study are provided in Supplementary Table 1.

All the MPXV genome datasets were aligned against the Clade II reference genome (which is an early hMPXV-1 genome from Nigeria, accession: NC_063383) using minimap2 v2.17 (*48*) to generate a single coordinate system. Genome sequences were trimmed from position 190,788 to the end, corresponding to the 3’ terminal repeat region of the MPXV genome. A series of repetitive or low-complexity regions were masked from the alignment (Supplementary Table 3). For convenience, this alignment and extraction pipeline is available as an installable tool at https://github.com/aineniamh/squirrel.

The full MPXV phylogeny of Clade I, IIa and IIb was estimated using maximum likelihood in IQ-TREE v2.0 using a Jukes-Cantor substitution model (*49*) and subsequently midpoint rooted. For the ancestral reconstruction analysis, we estimated phylogenies for each of Clade I, Clade IIa and Clade IIb. The MPXV Clade IIb phylogeny was estimated with the same parameters, but with the outgroup specified as the 1978 Nigerian MPXV genome sequence (Accession KJ642615). Similarly, we estimated phylogenies independently for the hMPXV-1 coding sequence alignment and for Clade I and Clade II of MPXV using the above parameters with genome sequence KJ642617 as an outgroup.

We performed the ancestral state reconstruction using IQ-TREE2 on all individual Clade phylogenies of MPXV. We parsed out all reconstructed sites that vary unambiguously across the phylogeny (i.e. we exclude missing data) and mapped the single nucleotide mutations that occurred to the relevant branch of the phylogeny. Using the reconstructed node states we also catalogued the dimer and heptamer context of all C->T or G->A mutations that occurred across the phylogenies. Scripts are available from http://github.org/hmpxv/apobec3/.

We compared each recent Clade IIb sequence with the reconstructed most recent common ancestor of the Clade IIb phylogeny (Figure 4) and collating the mutations down each branch from root-to-tip and their dinucleotide context. Using only APOBEC3-type mutations, we constructed an APOBEC3 root-to-tip linear regression for Clade IIb and similarly for lineage B.1 (phylogeny in Supplementary Figure 1). We performed a linear regression of the number of APOBEC3 mutations from the root of the tree to each tip against the date of collection of the tip. The slope is an estimate of the APOBEC3-specific mutation rate and assuming that APOBEC3 mutations are characteristic of infection in the human host, we use the X-intercept of this regression to estimate the date of emergence into a human host. For more details and the code, see Supplementary Info. This approach does not adjust for phylogenetic correlation and, in particular, the upward shift in B.1 is the result of a single shared branch with a higher than average number of mutations, but one that is not necessarily beyond expectations.

To investigate the target site context of mutations occurring in the hMPXV-1 phylogeny, we extracted the relevant nucleotide heptamers for all C->T or G->A mutations that occurred across the MPXV Clade IIb phylogeny (260 of 296 mutations) and for G->A mutations we took the reverse complement of the nucleotide sequences to normalise for strand. We also calculated the heptamer context for just the backbone of the phylogeny (i.e. the branches leading to the 2022 outbreak), which included 55 of 57 mutations on the respective branches. Using the reconstructed mutations and branch states in the phylogeny, we collected the ancestral dimer context for mutations that were either G->A or C->T to assess which mutation occurred in a context consistent with APOBEC3 editing.

## Results

The diversity of MPXV can be decomposed into three major clades; Clades I, IIa and IIb (Figure 1A; Happi et al., (*15*)). Clade I represents MPXV sampled in Central Africa and Clade IIa is composed of viruses from human and non-human animal samples taken in or connected to West Africa. Both of these clades include virus genomes spanning from the 1970s to present day. Clade IIb has an early sample taken in 1971 (KJ642617) but most of the sequences in Clade IIb are more recent virus genomes from 2017-2022 that (*15*) have labelled as hMPXV-1 (Figure 1A). Within the recent diversity of Clade IIb (indicated as the darker box within IIb in Figure 1A), we catalogued transmitted mutations that occurred between 2017 and 2022 with only a single representative from the 2022 global lineage B.1. The great majority of these mutations are of the type G->A or C->T (90.8%) (Figure 1B). Strikingly, the heptamers of C->T and G->A mutations that occurred across the Clade IIb phylogeny show this is a specific enrichment of APOBEC3-type dimer mutations of the type TC->TT and GA->AA (Figure 1C). Similarly, this enrichment of TC->TT and GA->AA mutations is observed within the B.1 lineage (Supplementary Figure 2).

Comparing MPXV Clade IIb with Clade Ia and IIa emphasises that this pattern is not seen outside of Clade IIb (Figure 1D-G). For the other MPXV clades, APOBEC-type mutations are observed at between 8 and 13% frequency, which fits with the expected proportion under standard models of nucleotide evolution (*50, 51*) (Figure 1D & F). Observed heptamers around the observed C->T and G->A mutations show a striking enrichment in TC and GA target sites in the genomes sampled from 2017-2022 in contrast with the rest of MPXV diversity (Figure 1E & G).

Considering all GA and TC dimer sites in the Clade II reference genome (Accession number NC_063383) – i.e. those that could be the target of APOBEC3 editing but had not been by that point – we looked at what the effect of a deamination mutation at these sites would be in terms of amino acid changes (Figure 4). Of the 23,718 such dimers, 61.6% (14,618) would produce amino acid replacements, 21% (4990) would be synonymous, 2.9% (692) would induce stop codons, and 14.4% (3418) would occur outside of coding regions. For the Clade IIb genomes, of the 633 mutations at these dimers that did occur, 38.7% (245) were amino acid replacements and 35.7% (226) were synonymous, 4.7% (30) were nonsense, and a further 132 APOBEC3 mutations were in intergenic regions (20.8%). The probability of getting 226 or greater synonymous mutations out of 663 under a simple binomial distribution with 21.0% chance of a context being synonymous is P=7.6×10^-18^. We do not see the same enrichment for synonymous mutations in the mutations that are not APOBEC3-like, although the quantity of these mutations is considerably lower (Supplementary Figure 3). There are also more mutations outside of protein coding regions than we would expect based on the location of target dimers (probability of 4.5×10^-6^ of getting at least 132 non-coding mutations given only 14.4% of targets are in these regions). This supports the hypothesis that what we are observing are the residual least harmful APOBEC3 mutations after natural selection has eliminated those with substantial fitness costs to the virus. By comparing the density of observed C->T and G->A mutations and the density of APOBEC3 target sites (TC or GA dinucleotides) across the reference genome, we see that the distribution of mutations is not merely a product of the availability of target sites (Kolgorov-Smirnov test statistic=0.07, P-value=0.0002; Figure 2B-C). When considering synonymous and non-synonymous APOBEC3-like mutations separately, we observe the difference between target and observed APOBEC3-like mutations for both cases (Supplementary Figure 4).

The ‘repertoire’ of mutations that APOBEC3 is able to provide as genetic variation on which natural selection can act is severely restricted. Only a limited number of dinucleotide contexts are present, and the amino acid changes that APOBEC3 editing can induce is also limited (Figure 3). Only 13 different amino acid replacements are possible, and three that give rise to stop codons, and they are not reversible by the same mechanism. This means that given the restricted set of positions at which these mutations occur and the limited amino acid changes they can result in, we don’t expect the elevated rate to change the rate of adaptation of the virus substantially.

Since 2017 the genomes thus far sampled from Clade IIb have accumulated APOBEC3-type single nucleotide mutations approximately linearly over time (Figure 4b). The estimated rate of accumulation was 6.18 per year (95% credible intervals of 5.20, 7.16). For the B.1 lineage the rate was 5.93 per year (2.95, 8.92) – suggesting that despite widespread and rapid transmission within MSM networks, the rate of accumulation of APOBEC3 mutations remained the same as the rest of the Clade IIb. It is notable that the regression line for B.1 lies substantially above that for the rest of Clade IIb suggesting that this lineage accumulated more mutations than expected prior to the emergence of B.1. However most of these mutations are also present in the genome from Maryland, USA from November 2021 (Accession number: ON676708), indicating they arose and circulated for some months prior to the B.1 epidemic.

Extrapolating back to number-of-suspected-APOBEC3-mutations (y-axis) = 0 provides an estimate of when the first APOBEC3 mutations occurred in the stem of the branch leading to the 2017 epidemic. We estimate this to be 20-Jun-2015 (95% credible intervals of 6-Oct-2014, 3-Mar-2016) which represents the likely time range in which the initial jump to sustained human to human transmission occurred.

## Discussion

Within MPXV Clade IIb, we observe rates of molecular evolution far greater than that expected for double-stranded DNA viruses and indeed that observed in Clades I and IIa of MPXV (*20*). The recent diversity in Clade IIb overwhelmingly consists of TC->TT and GA->AA nucleotide changes, consistent with APOBEC3 enzyme editing, and a recent study has demonstrated APOBEC3F editing during human MPXV infection (*52*). Since 2022, the B.1 lineage has been sampled and sequenced internationally in the global epidemic of MPXV. Lineage B.1 is known to be circulating by sustained human-to-human transmission and as such, mutations that have accumulated in B.1 can be considered characteristic of this mode of transmission. Our analysis highlights that evolution within Clade IIb prior to the emergence of lineage B.1 mirrors that within lineage B.1, but is distinct from MPXV Clade I or IIa.

We suggest that the APOBEC3-driven evolution of recent Clade IIb MPXV is a signature of a switch to sustained transmission within the human population. Within the B.1 lineage, believed to be entirely the result of human infection and transmission we continue to see the same pattern of predominantly APOBEC3 mutations accumulating at a similar rate to that seen in A lineage genomes since 2017. It is unlikely that, by chance, MPXV evolved to become susceptible to APOBEC3 action within the putative rodent reservoir prior to the emergence of cases and to retain that susceptibility to human APOBEC3 molecules once transmitting in humans. Given that all human cases sequenced since 2017 share substantial numbers of APOBEC3 mutations, including nine on the stem branch leading to hMPXV-1 it is very unlikely these represent multiple zoonotic introductions.

If we assume this observed evolution within hMPXV-1 is APOBEC3-driven, this may have implications for its sustained transmission in the human population. APOBEC3 hypermutation is a host-mediated antiviral mechanism. These molecules act as the viral genome is being replicated and single strands are exposed. During repeated rounds of replication either strand can be deaminated leading to both C->T and G->A changes on the positive strand as seen here. Thus it is likely that the genomes that are extensively mutated by APOBEC3 will simply not be viable and will not be transmitted further. The extensive rounds of genome replication that MPXV undertakes in the cytoplasm likely means that most genomes are not affected by APOBEC3 action. Occasionally however a genome, only modestly mutated by APOBEC3, may remain viable and be transmitted. We see this in the enrichment of observed synonymous and intergenic mutations relative to available targets in the MPXV genome. Given the non-reversibility of the APOBEC3 action, sustained evolution within the human population may result in a depletion of lower-consequence target sites (i.e., synonymous or conservative amino acid changes) and thus expedite a decrease in fitness of MPXV over time. This could be both through a reduction in the number of viable offspring viruses produced by infected cells and as a result the accumulation of moderately deleterious mutations by genetic drift. However the timescale on which this might happen is uncertain and other evolutionary forces such as recombination may act to restore fitness. A further uncertainty arises from the number of genes associated with host immune modulation in poxvirus genomes. Mutations that alter or abrogate the function of these genes may have little direct effect on virus replication machinery, but may disrupt the virus/host interaction. There is precedent for the naturally occurring inactivation of such genes in VARV, and consequently the loss of function of some MPXV genes through APOBEC3 activity may potentially have adaptive value for the virus (*53, 54*).

Even if the mutations that accumulate through this process are simply the neutral residue of a sub-optimal antiviral host defence, they have produced sufficient variability for the phylogenetic analysis of the epidemic over the short term. The initial lineages proposed by Happi et al (*15*) have now expanded with the 2022 epidemic B.1 lineage now encompassing 12 sublineages [github.com/mpxv-lineages/lineage-designation]. The rapid and temporally linear accumulation of mutations means that genomic epidemiological models and tools (*55, 56*), usually employed for RNA viruses, are also applicable to hMPXV-1.

Since the identification of the B.1 lineage, a number of countries have reported other lineages that lie outside the diversity of B.1, including USA, UK, Portugal, India and Thailand. In almost all instances these cases are reported as having a history of international travel. All of these lineages (designated as A.2.1, A.2.2, A.2.3 and A.3) can be phylogenetically traced back to the epidemic in Nigeria (Figure 4a). This suggests that sustained human to human transmission is still ongoing outside of recognised MSM networks and stopping transmission in the MSM communities, whilst necessary, will not be sufficient to eliminate the virus as a human epidemic. There are large portions of the globe without the surveillance to detect MPXV cases and if sustained human to human transmission has been ongoing since 2015-2016 it is highly plausible there are other populations that are currently enduring epidemics.

Historically, MPX was considered a zoonotic disease and cases have been treated as independent spillover events with low levels of circulation in the human population. This assumption has informed outbreak control measures as low levels of spillover at the human/reservoir interface would suggest more targeted interventions were necessary rather than a large-scale vaccination campaign. In the light of our observations, this historic assumption may no longer be valid when considering MPX cases and future outbreak control measures should treat any novel zoonotic outbreak of MPXV as having the potential for sustained human-to-human transmission, regardless of whether infections are spreading internationally or within a country that has an endemic reservoir host.

---

**Supplementary Figure 1.**
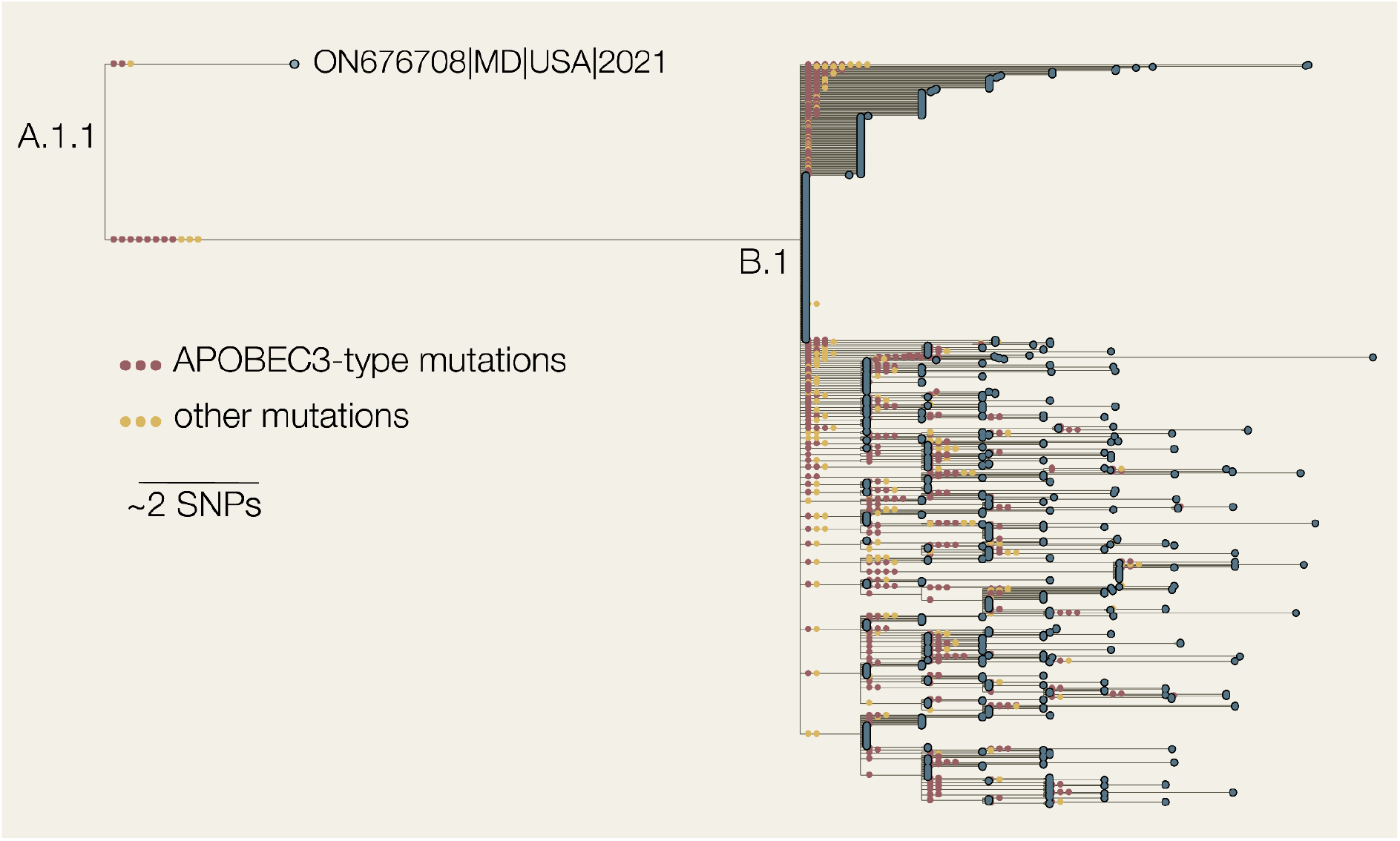
Phylogeny containing all high-quality hMPXV1 B.1 genomes shared on Genbank as of 2021-08-22, with the 2021 Maryland sample (ON676708) used as an outgroup (n=769).

**Supplementary Figure 2.**
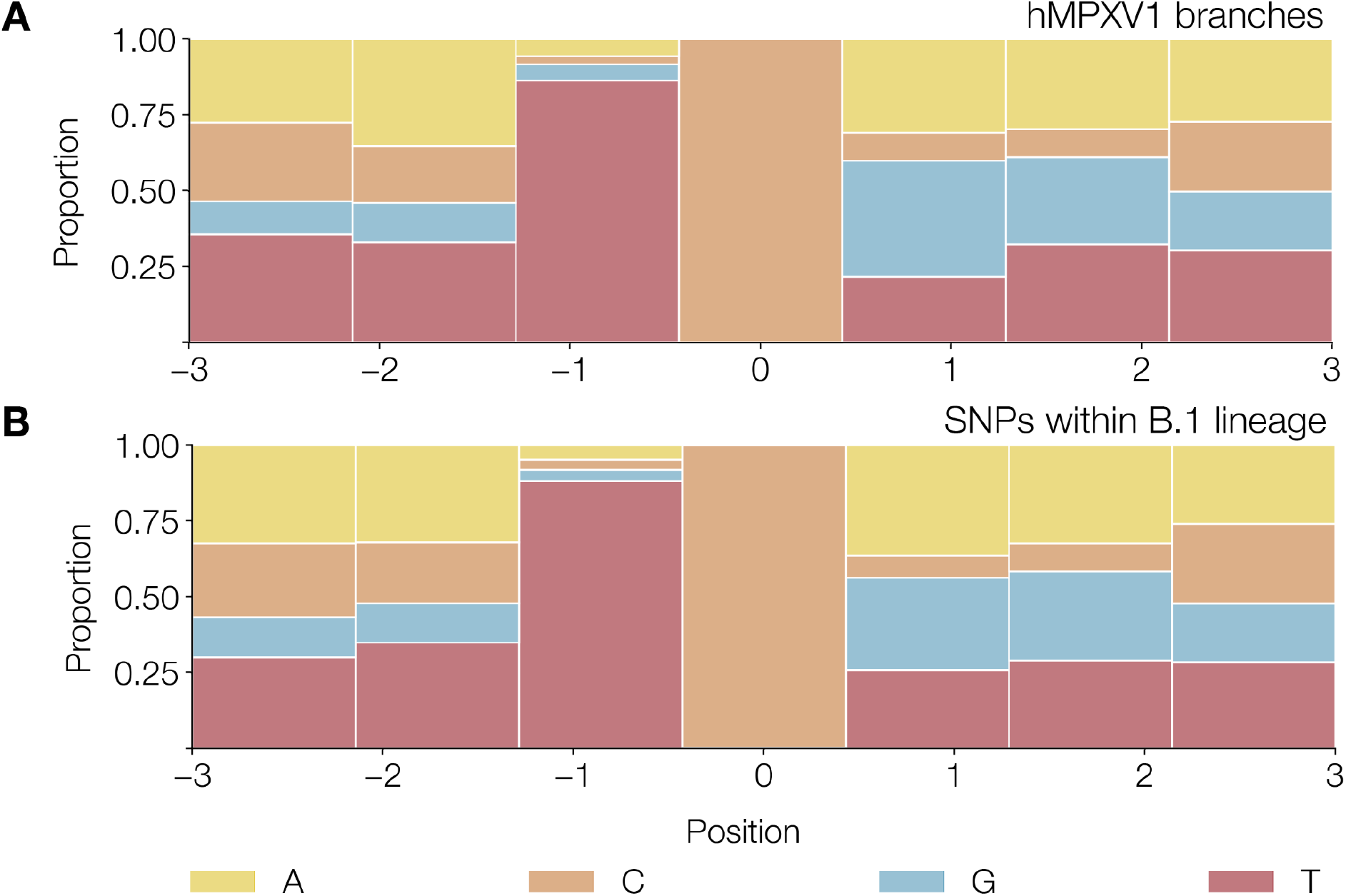
A. Observed heptamers of C->T or G->A mutated sites in 337 of 383 unambiguous mutations that occur along branches in the MPXV phylogeny in Figure 4A, excluding the mutations that occur on the 1971 branch. Heptamers associated with G->A mutations have been reverse-complemented to reflect deamination on the negative strand. B. Observed heptamers of C->T or G->A mutations in the B.1 lineage of Clade IIb (677 of 836 mutations), which we know is being transmitted human-to-human.

**Supplementary Figure 3.**
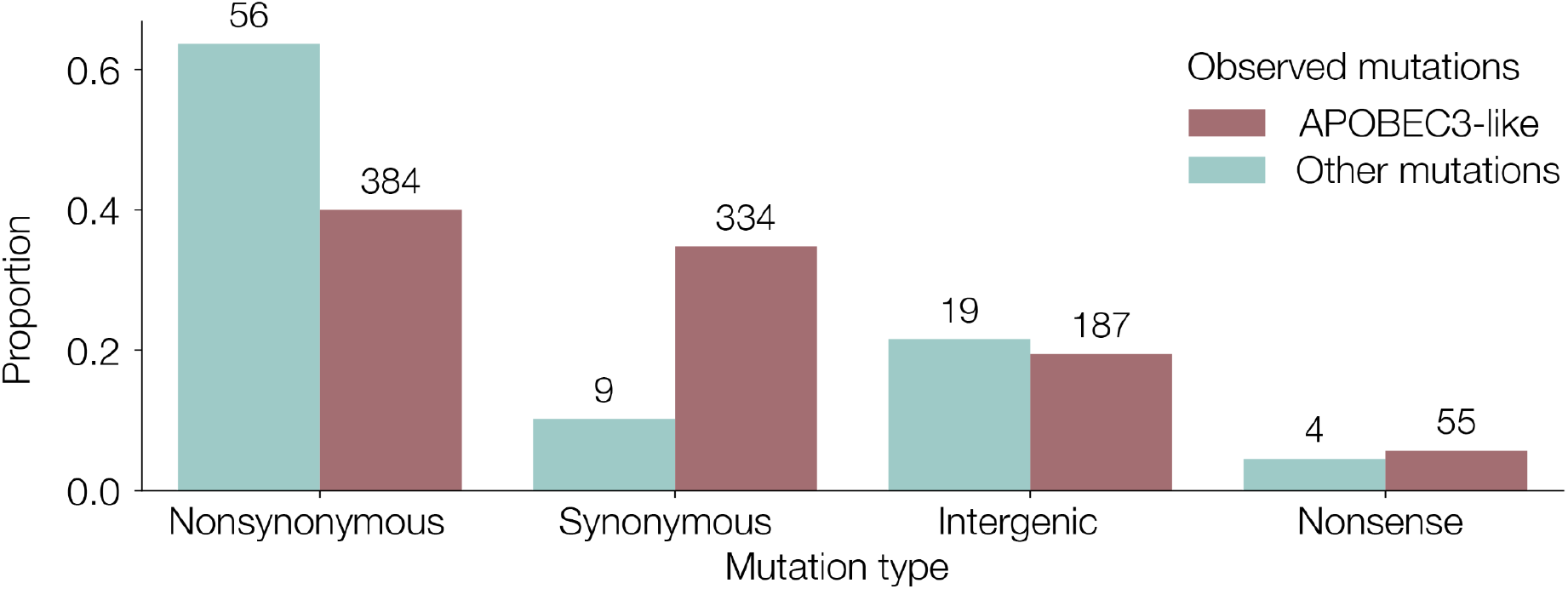
Observed mutations across the Clade IIb phylogeny (including genomes in Figure 4 and Supplementary Figure 1, except the outgroup branch leading to the 1971 genome sequence). These are categorised into non-synonymous (altered amino acid), synonymous (amino acid remaining unchanged), nonsense (editing producing a stop codon) and intergenic (not present in a coding sequence), and coloured by whether they have occurred in an APOBEC3-like context or not.

**Supplementary Figure 4.**
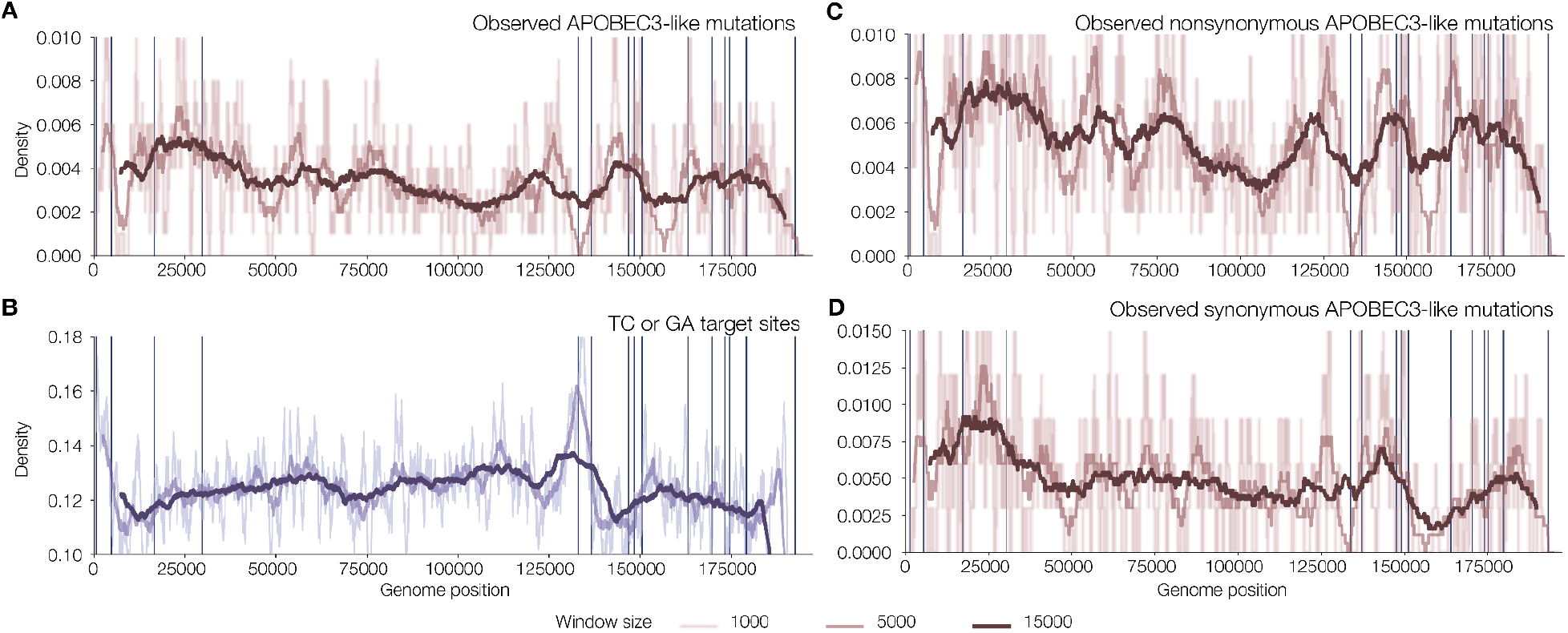
The density (moving average linear convolution with window sizes of 1000, 2500 and 15000) of A) observed APOBEC-like mutations across the MPXV genome, B) available APOBEC3 target sites in the Clade IIb reference genome (Accession NC_063383), C) observed APOBEC-like mutations that result in synonymous changes and D) observed APOBEC-like mutations that result in nonsynonymous changes. We see the same difference between target density for both observed synonymous and non-synonymous distributions (Kolgorov-Smirnov test statistic synonymous vs target = 0.1133, P-value=0.0003; Kolgorov-Smirnov test statistic non-synonymous vs target = 0.067, P-value=0.059). Repetitive masked regions indicated by vertical bars.

**Supplementary Table 1:**
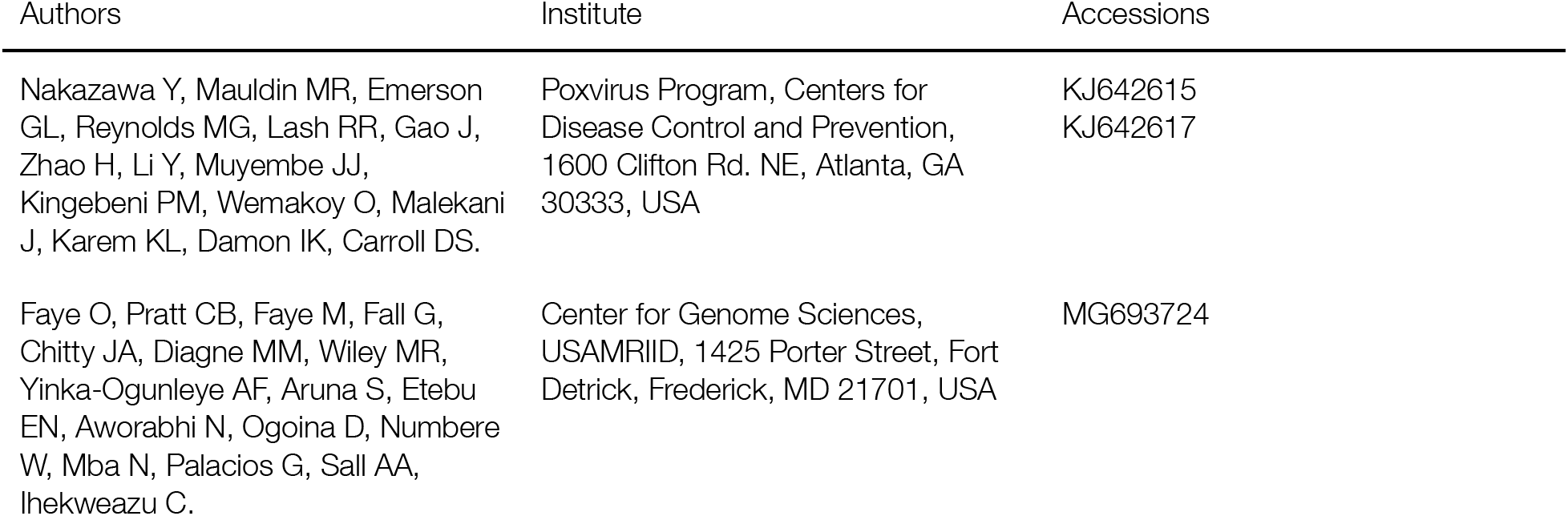
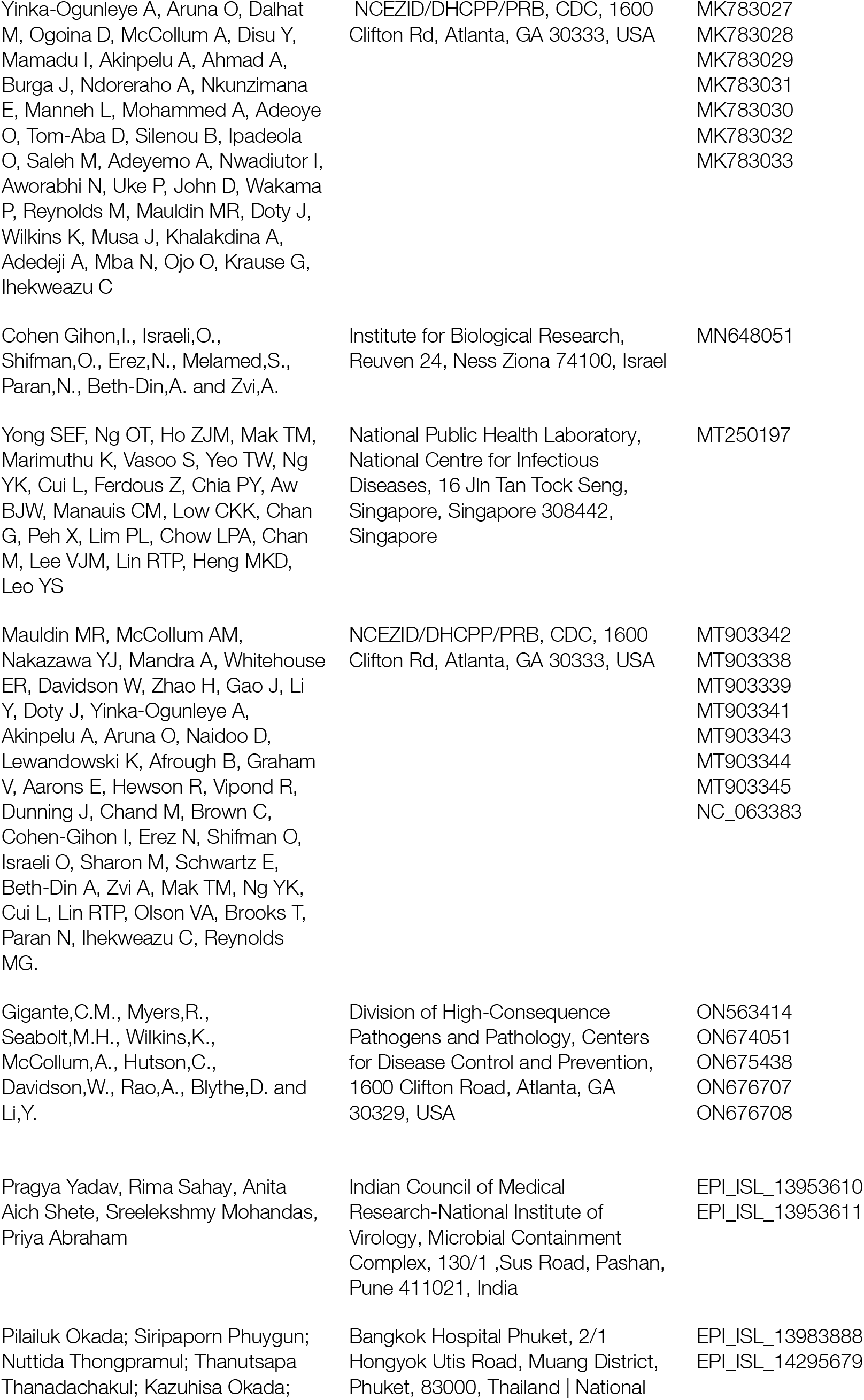
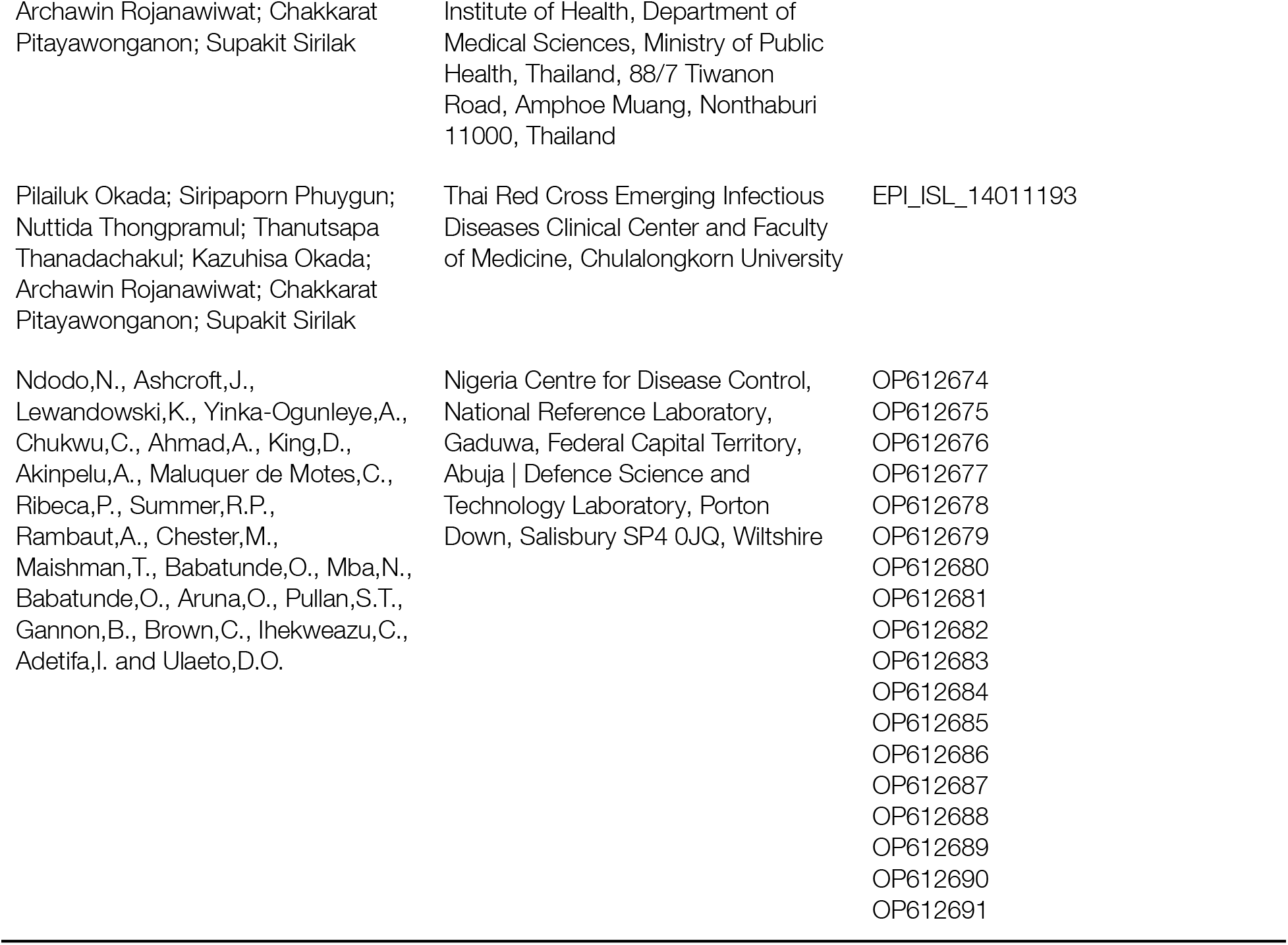
Authors, Institute and Accession numbers for sequences in analysis of hMPXV1.

**Supplementary Table 2:**
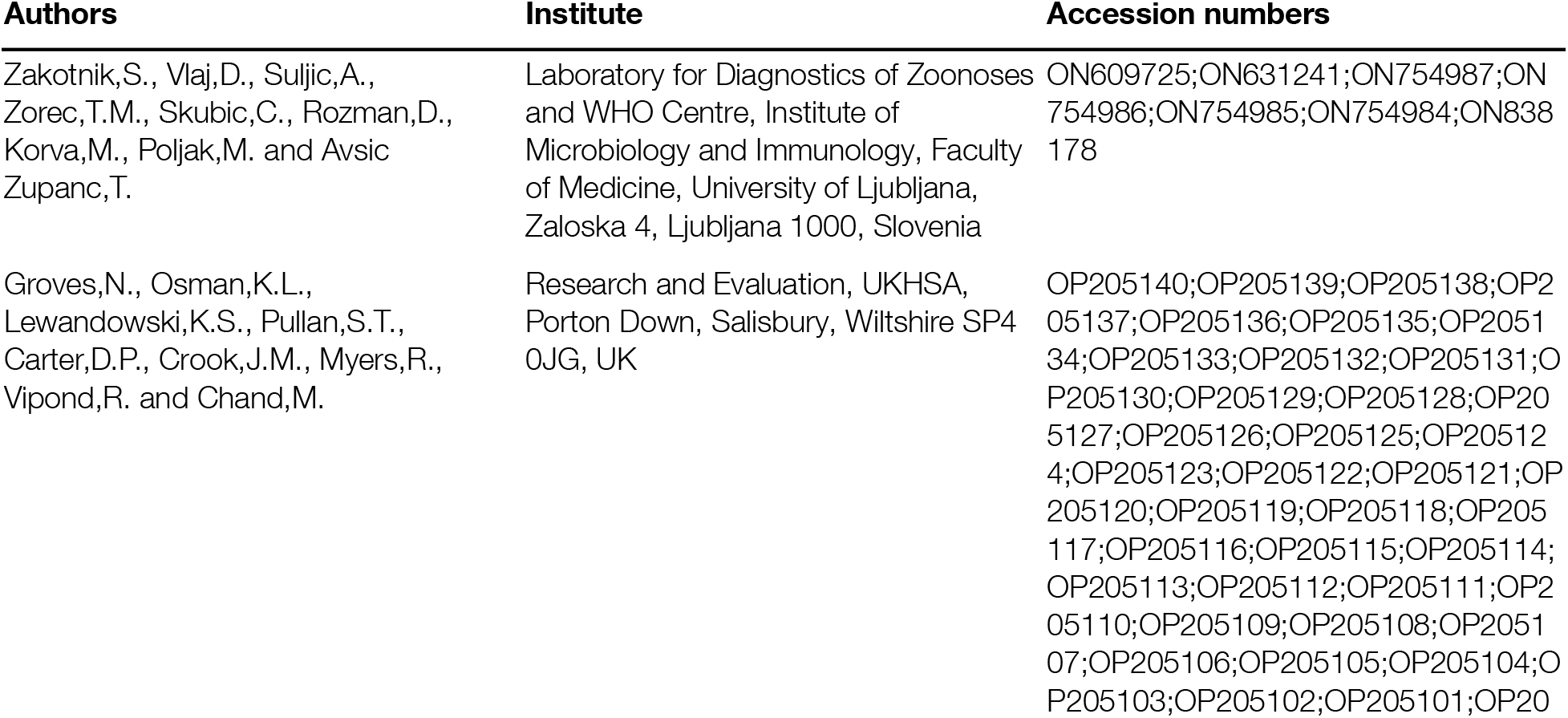
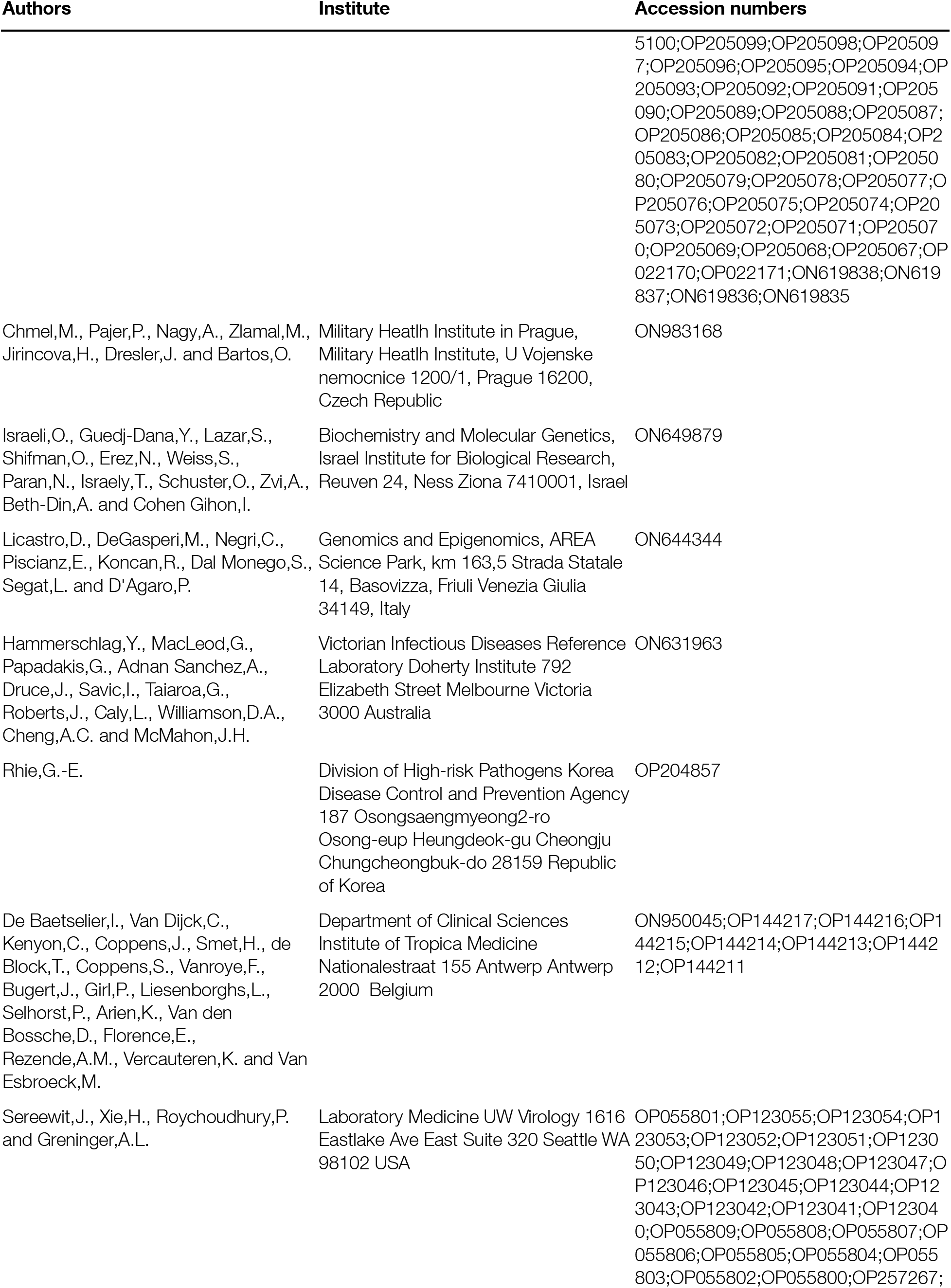
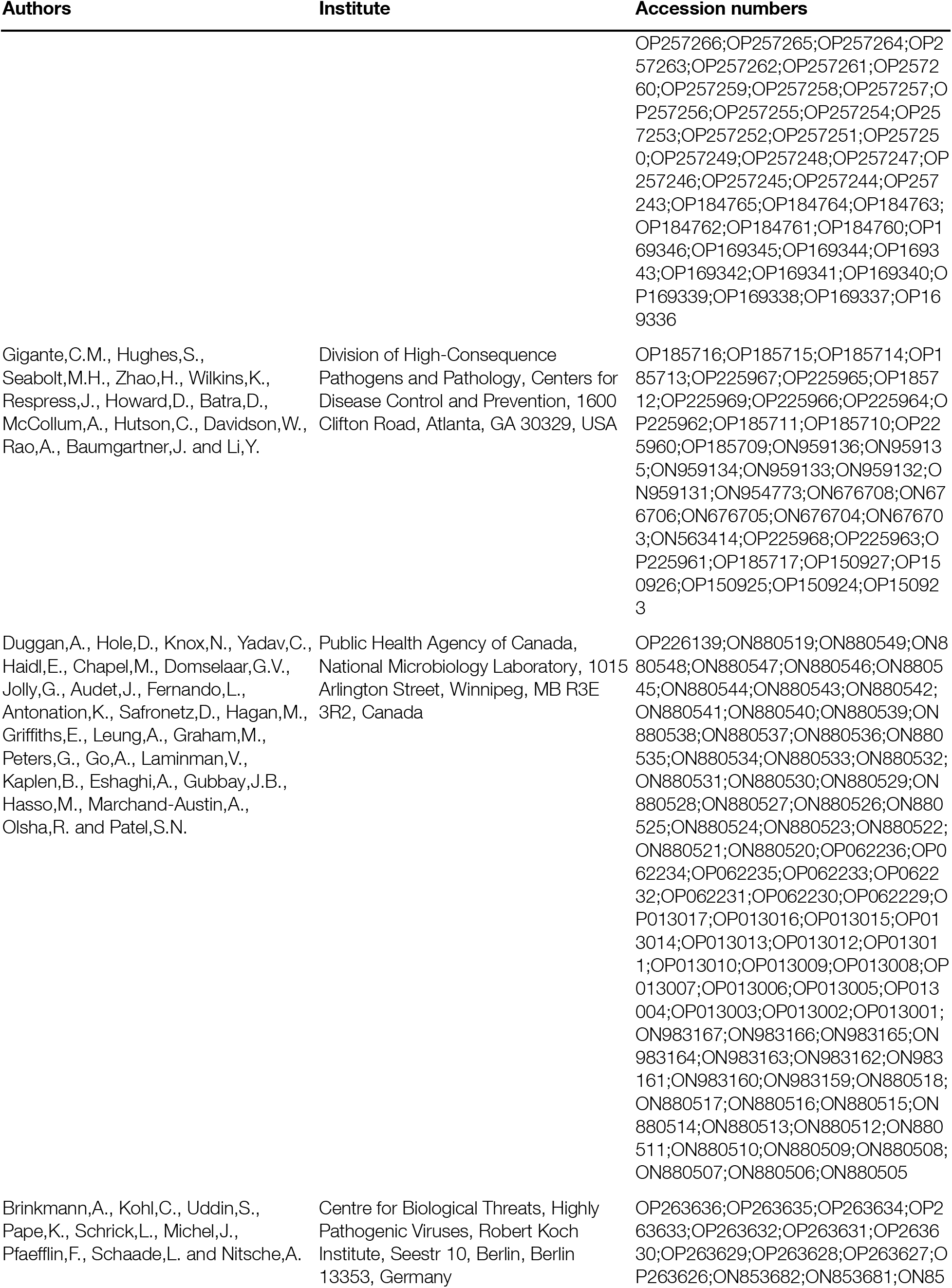
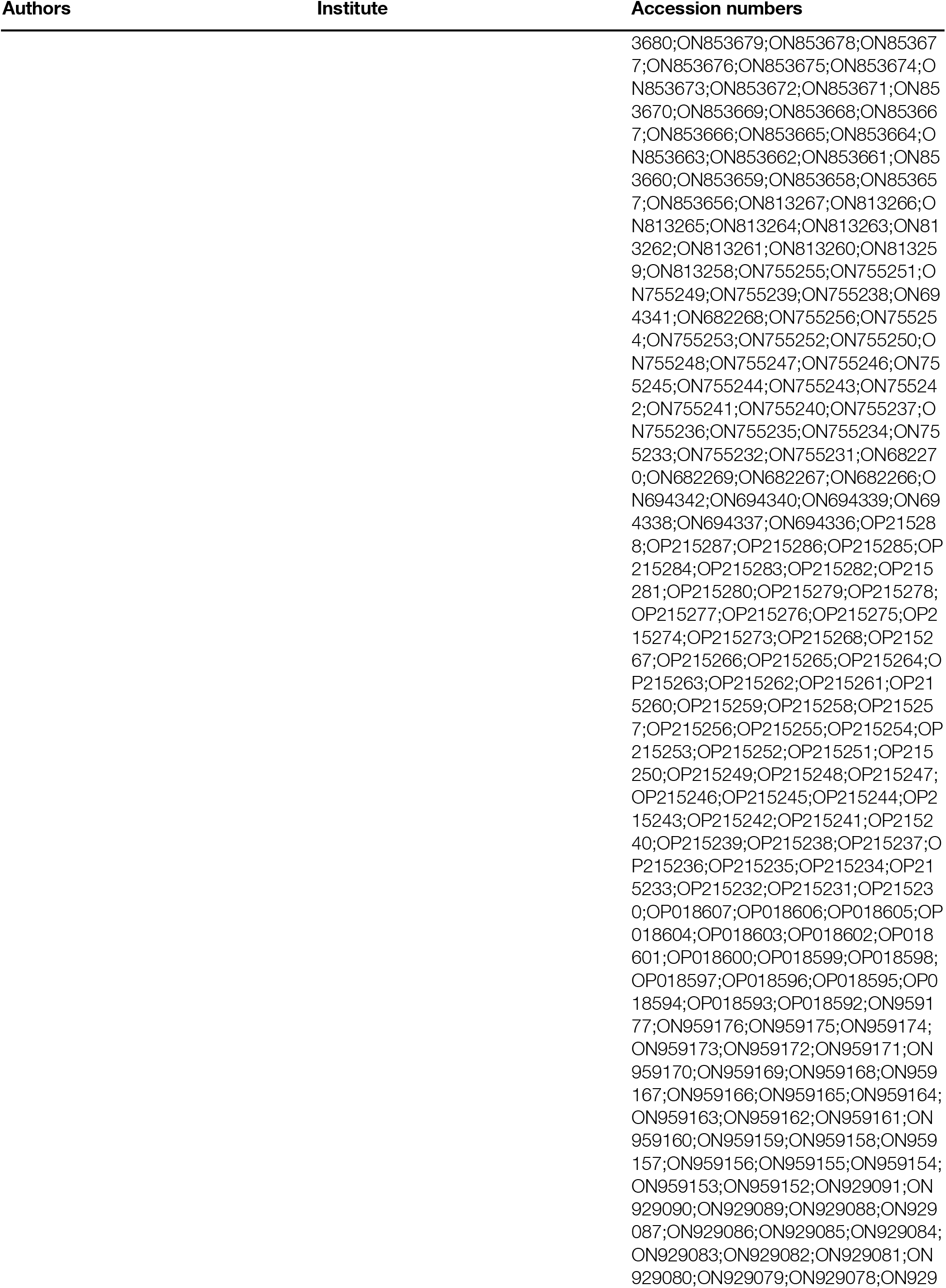
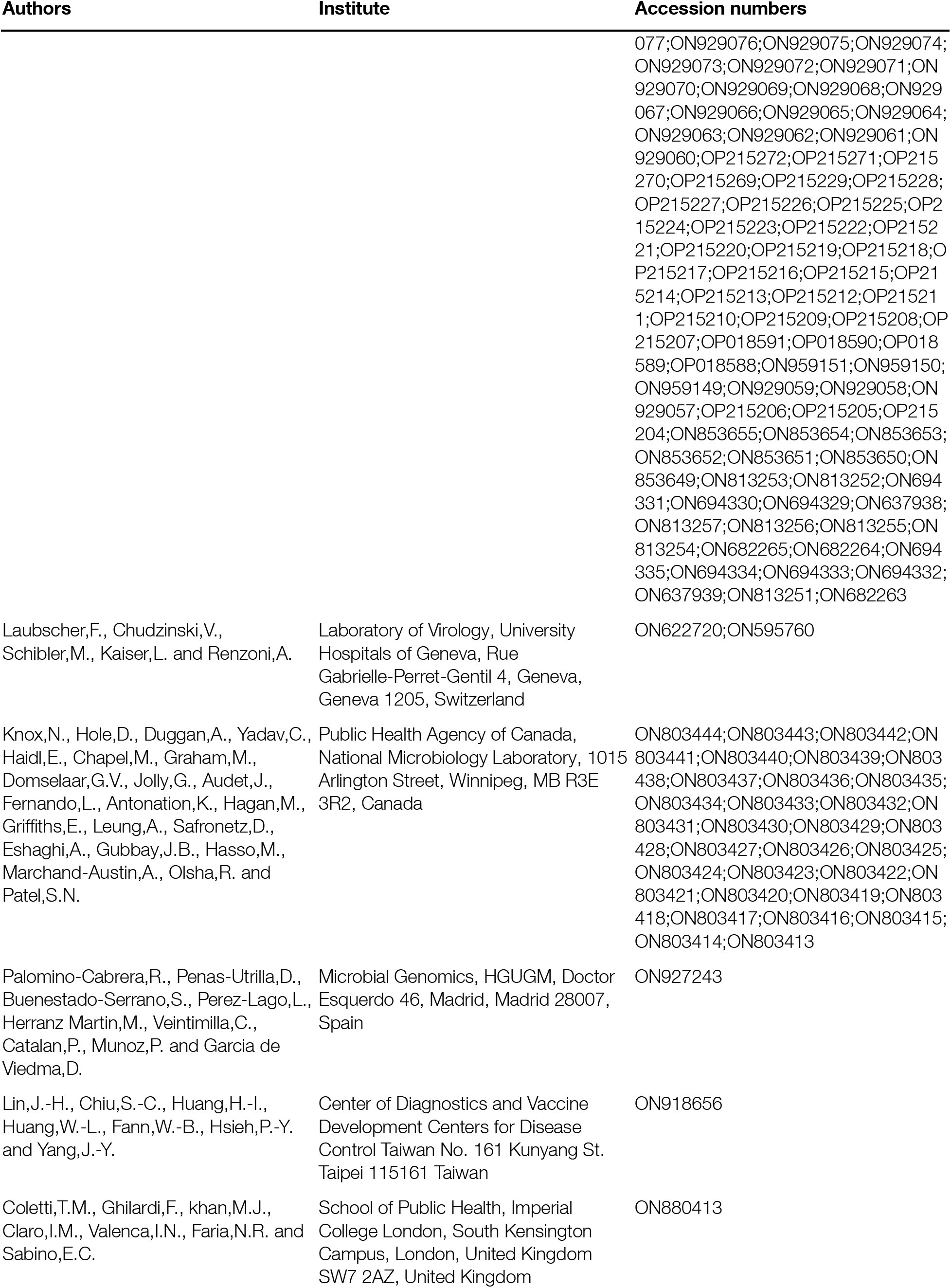
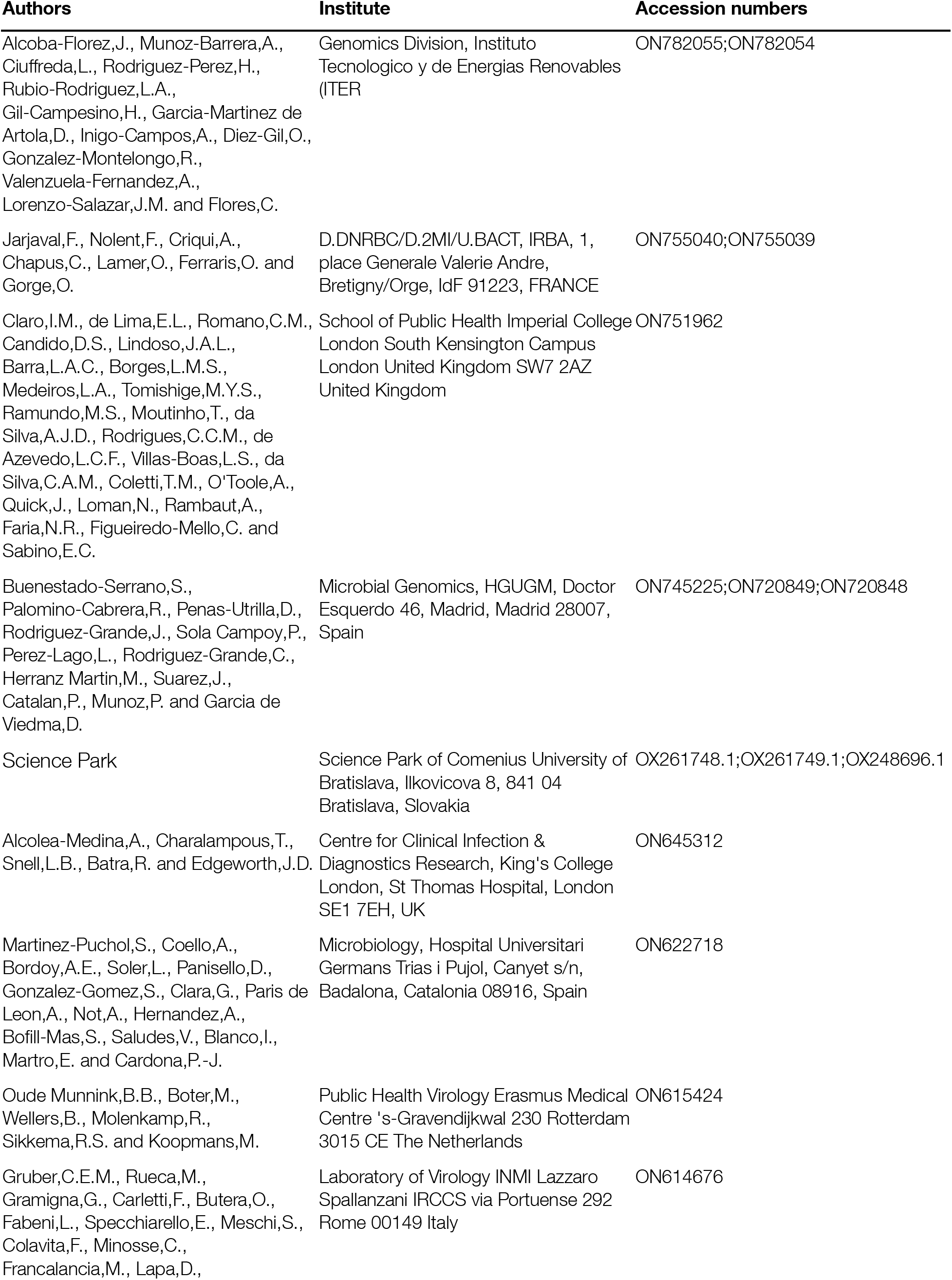
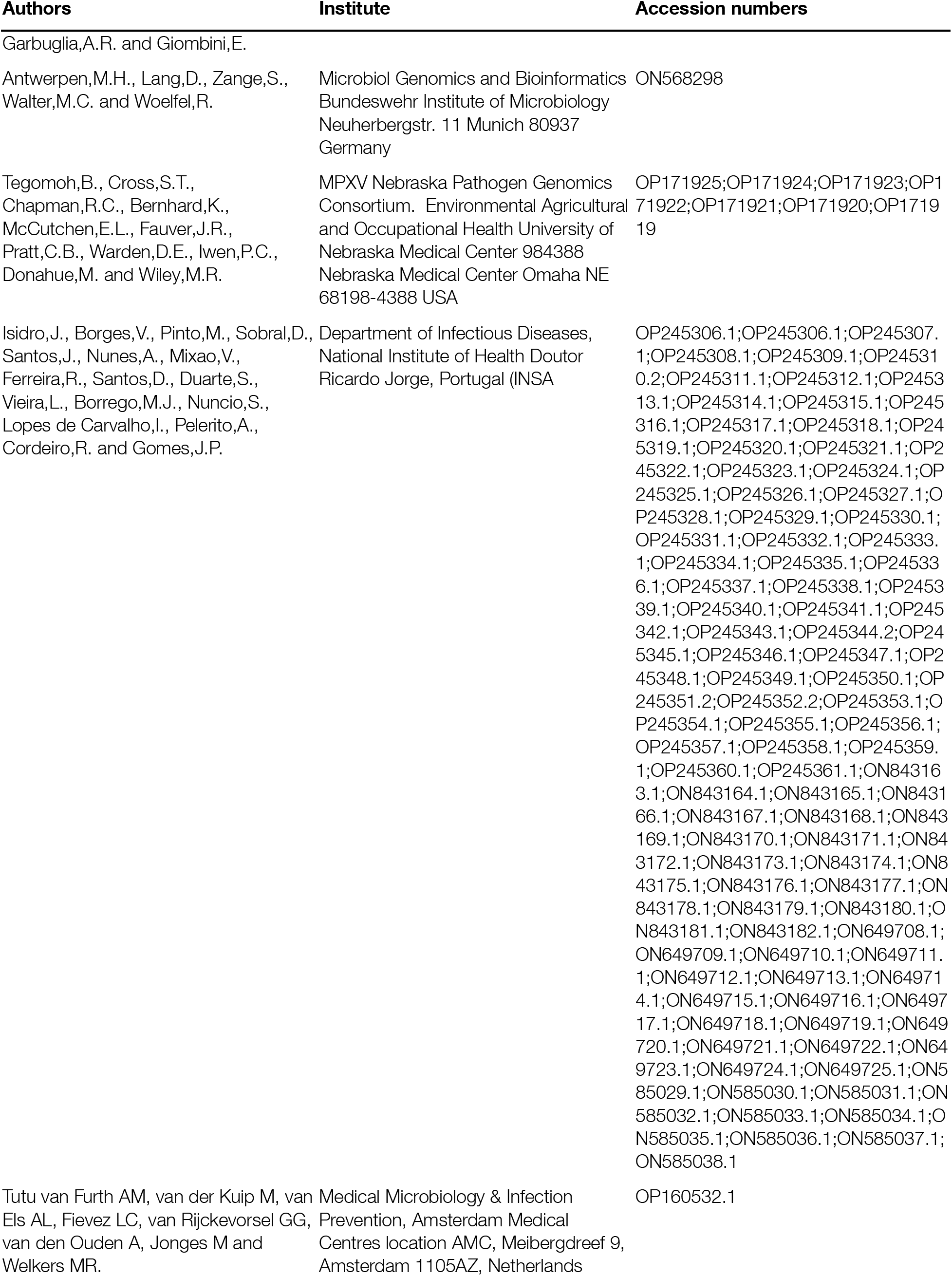
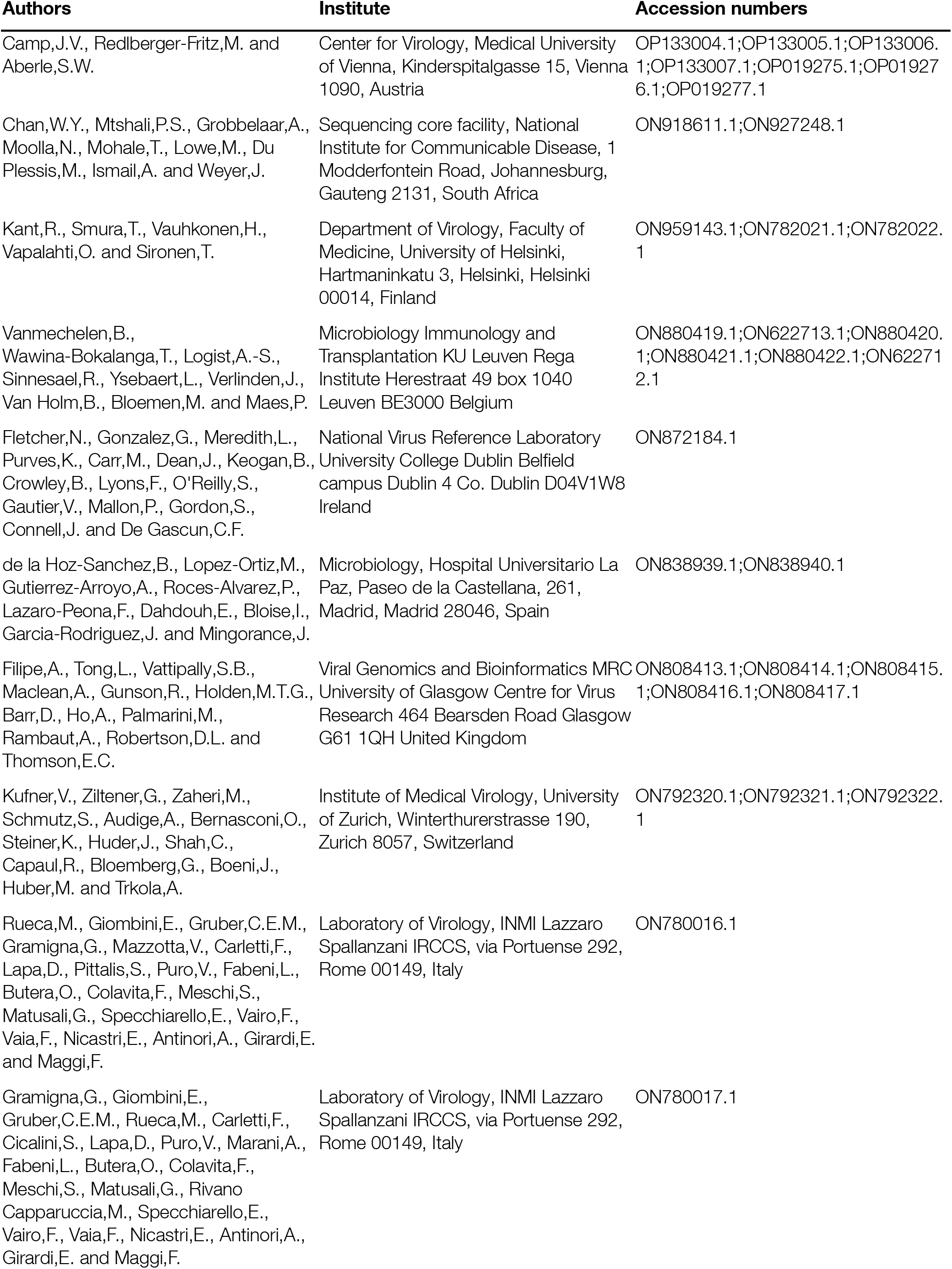
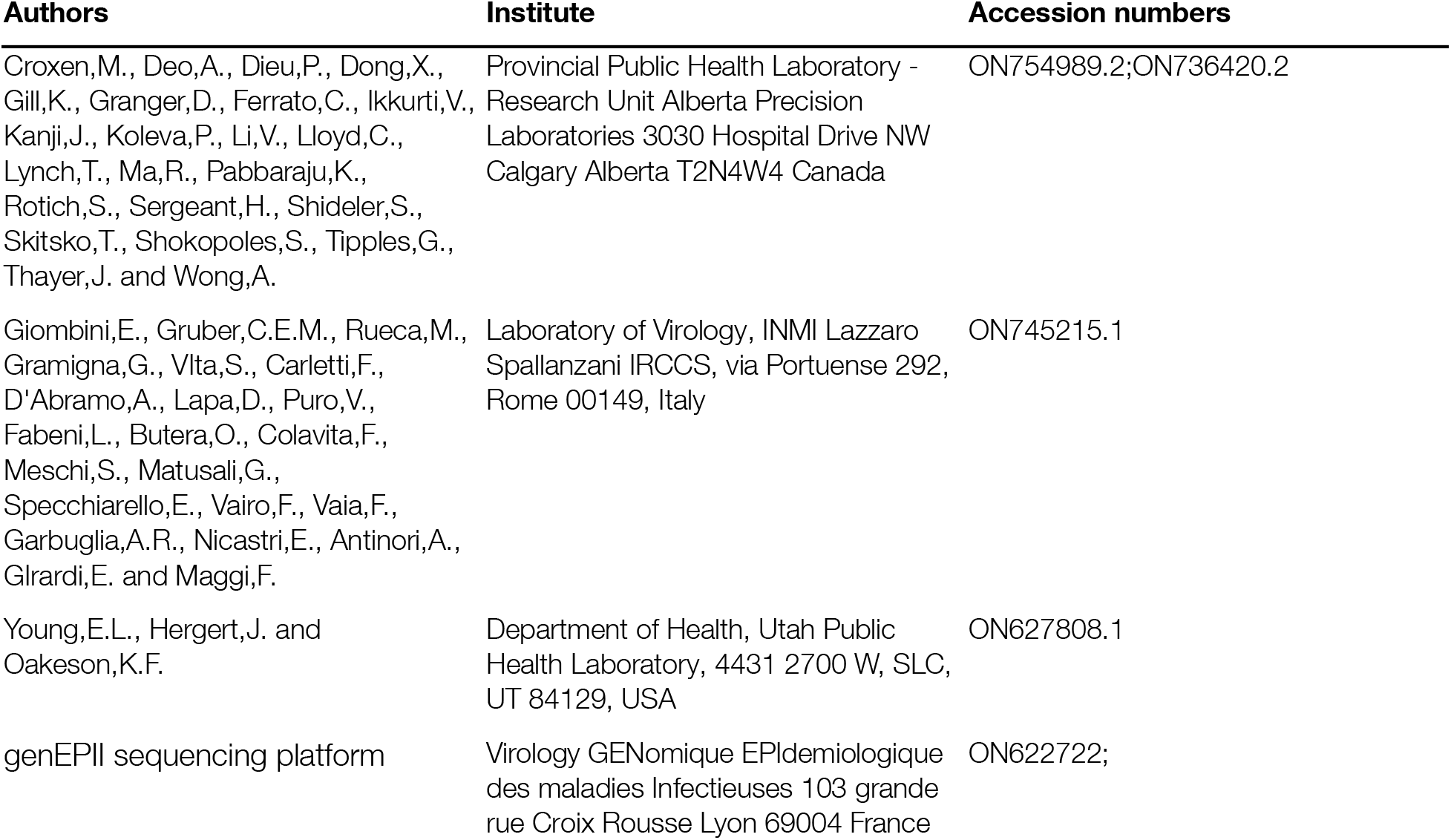
Authors, institute and accession numbers for sequences in Supplementary Figure 1.

**Supplementary Table 3:**
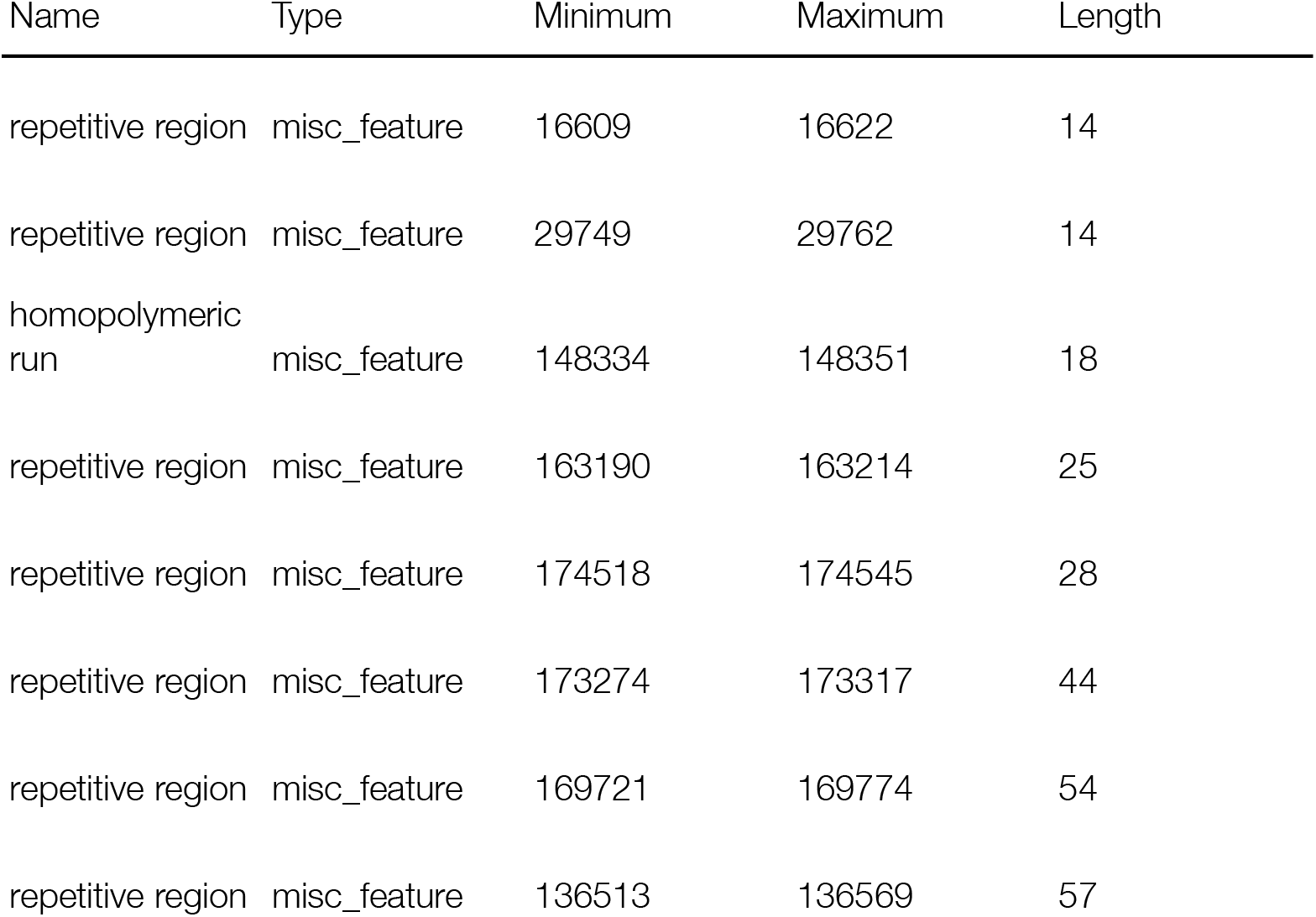
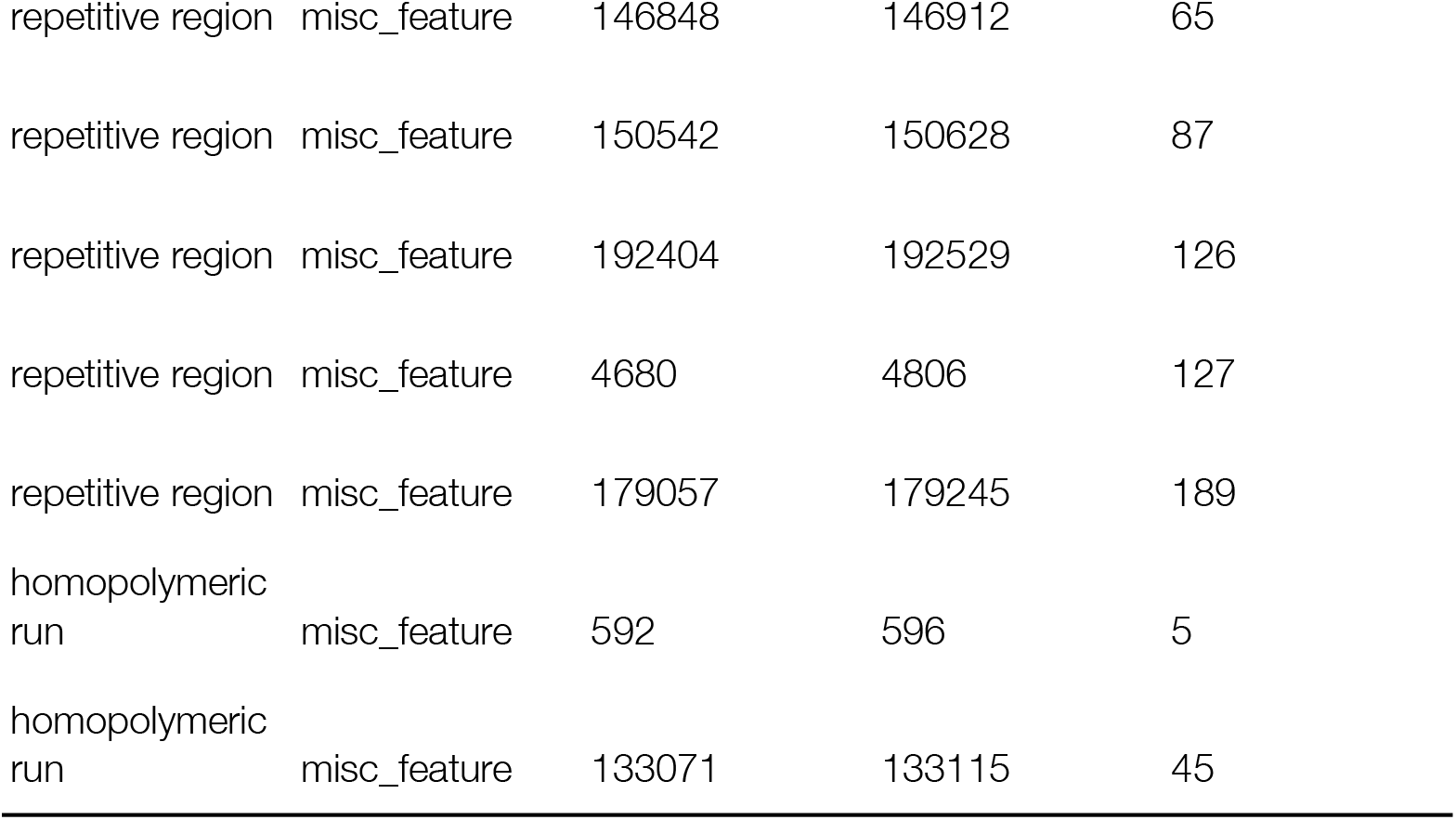
Regions masked in alignment.

## Supplementary Methods

The model for the linear regression of number of APOBEC3-type mutations against time was as follows:

m_i_ ~ Normal(μ, σ)
μ = α + β (*t*_i_ - *t*_mean_)
α ~ Normal(11, 100)
β ~ Lognormal(0, 1) and
σ ~ Normal(0, 20),

where m_i_ is the number of mutations for genome i,

*α* is the y-intercept and has a prior centred on the minimum number of mutations observed over all genomes, *β* is a strictly-positive evolutionary rate per year, *t*_i_ is the time of collection of the sample and *σ* is the model error standard deviation.

The model was fitted to the data in a Bayesian framework using the quadratic approximation implemented in the *rethinking* package in R (McElreath, 2020). Posterior estimates and 97% highest posterior densities of the parameters for the lineage A data are *α*: 25.24 (23.80, 26.66), *β*: 6.18 (5.19, 7.16), *σ*: 4.76 (3.74, 5.78) and for the lineage B.1 data are *α*: 57.67 (57.49, 57.85), *β*: 5.93 (2.95, 8.91), *σ*: 1.77 (1.64, 1.90). An R script performing this analysis and generating the graphic seen in Figure 4b is available at http://github.org/hmpxv/apobec3/

McElreath (2020). Statistical Rethinking: A Bayesian Course with Examples in R and Stan (2nd edition). CRC Press.

